# Mechanical tension expands the microtubule lattice stepwise and modulates kinesin-1 binding in an isoform-dependent manner

**DOI:** 10.64898/2026.06.17.732986

**Authors:** Yannic Lurz, Benedikt S. J. Fischer, Jaya Mishra, Laura Muras, Erik Schäffer, E. Michael Ostap, Nisha Mohd Rafiq, Igor M. Kulić, Serapion Pyrpassopoulos

## Abstract

Recent work has shown that the microtubule lattice possesses remarkable structural plasticity, with its conformation modulated by microtubule-associated proteins and motor proteins. However, how this plasticity responds to mechanical forces remains poorly understood. Here, we developed optical tweezers and fluorescence microscopy assays to measure the effect of tensile forces on single microtubules. Quantum dot decoration enabled nanometre-precision measurement of lattice distortions of ∼0.33% under a change of mean tensile force of ⟨*ΔF*⟩ = 10.6 pN, within the range F_min_ = 1.29 pN to F_max_ = 22.3 pN — comparable to forces from one to three kinesin-1 motors. Within this force range, the binding rate of kinesin-1 isoform KIF5B decreased reversibly within seconds by ∼20% and the dissociation rate increased by ∼10%, reducing mean run length, that in extreme cases decreased by up to 46%. Substantial heterogeneity was also observed along individual microtubules, where distinct lattice regions responded differently to applied force, implying that lattice expansion is not always uniform. Consistent heterogeneity was observed in cells, where MAPs with competing conformational preferences assembled in non-overlapping patches along the same microtubule. A cooperatively-switching lattice Ising model based on tubulin conformational bistability, supported by dynamics simulations, quantitatively reproduces these observations with a critical switching force F_c_ = 8.5 pN, similar to established mechanosensory proteins such as talin and αE-catenin. Strikingly, no significant effects were observed for KIF5C, revealing a kinesin isoform-dependent mechanoresponse. Together, these findings establish microtubules as mechanochemical signal transducers, converting mechanical forces into biochemical signals with the speed, spatial precision and sensitivity required for rapid cellular responses.

**Significance Statement:** Microtubules have been implicated as mechanotransducers in both mammalian and plant cells, yet a physical characterization of how mechanical forces are sensed and transduced into biochemical signals has been lacking. The present study demonstrates that modest tensile forces of less than 20 pN are sufficient to expand cooperatively the microtubule lattice by ∼0.3%, which in turn modulates its biochemical interactions with kinesin-1 in an isoform-dependent manner, selectively affecting KIF5B motor activity but not KIF5C. Strikingly, this mechanotransduction occurs on a timescale of seconds, implying that microtubules are highly efficient conduits for propagating mechanical information across the cell body. These findings establish microtubules as *bona fide* mechanochemical signal transducers with the speed and sensitivity required for rapid cellular responses.

## Introduction

Microtubules are polar cytoskeletal filaments important for critical process of eukaryotic cells such as cell division and intracellular trafficking. They are formed by the GTP-dependent, non-covalent polymerization of αβ-tubulin heterodimers into linear, polar protofilaments, which interact laterally to generate the characteristic hollow cylindrical architecture of the microtubule (1, 2). Incorporation of αβ-tubulin into the microtubule lattice triggers the GTPase activity of αβ-tubulin, such that mature microtubules consist of a GDP-tubulin shaft capped by a region of GTP-bound tubulin at the growing plus end (3, 4). GDP lattice is known to adopt a compacted and unstable conformation while the GTP lattice is stable and expanded (5–8) and acts as protective cap preventing microtubule depolymerization (9). Loss of the stabilizing GTP cap, exposes the GDP-tubulin lattice and promotes rapid depolymerization as GDP-bound tubulin relaxes toward its preferred curved conformation breaking the lateral contacts between the protofilaments (9–11). As a consequence of the nucleotide dependent conformations of αβ-tubulin, microtubules are highly dynamic polymers that undergo repeated phases of growth and rapid depolymerization, termed catastrophe, both *in vivo* and in vitro (9, 12–14).

Beyond the nucleotide-dependent conformational switch, it has long been proposed that polymerised tubulin can access additional conformational states independently of its nucleotide state, a behaviour referred to as structural plasticity (15–17). Consistent with this idea, structural studies have shown that the conformation of the microtubule lattice can undergo dynamic, asymmetric fluctuations deviating from helical symmetry (18) and that its conformation can be further modulated by interactions with microtubule-associated proteins (MAPs) and motor proteins. At saturating concentrations, different MAPs and motors stabilise distinct lattice states, with tubulin adopting a spectrum of compacted or expanded conformations characterised by varying degrees of lattice twist depending on the binding partner (6, 19–23). Intriguingly, despite being GDP-bound, axonal microtubules in neurons were recently found to adopt an expanded, GTP-like lattice conformation rather than the compacted state expected for GDP-tubulin (24). Moreover, at sub-saturating concentrations, structural perturbations induced locally by kinesin motors are thought to propagate along the lattice and be sensed over distances of several micrometres by neighbouring motors, implying long-range allosteric communication through the microtubule lattice (25, 26).

Mechanical forces applied along the microtubule axis have also been shown to influence interactions with MAPs and motors. In particular, tensile forces ≤10 pN generated by a single kinesin-1 molecule modulate its detachment kinetics from microtubules over more than an order of magnitude (≈0.26–10 s⁻¹ (27)), whereas substantially higher forces (∼40 pN) significantly increase the dissociation of tau from tau decorated microtubules in the absence of soluble tau (22). However, despite growing evidence that both biochemical and mechanical cues tune microtubule structure, how microtubule structural plasticity responds quantitatively to applied mechanical tension remains poorly understood. Specifically, it remains unknown whether tensile forces within the range 10 – 20 pN, which can be generated by one to three kinesin-1 or mammalian cytoplasmic dynein molecules (27, 28), are sufficient to induce detectable lattice deformations, and if such mechanically induced changes influence the binding affinity of MAPs and motors at the single-molecule level.

Here, using dual- and single-beam optical trapping combined with high-contrast fluorescence imaging, we directly quantify longitudinal microtubule lattice expansion under physiologically relevant tensile forces. We find that modest increases in tension (∼10 pN) induce a measurable and, on average, reversible expansion of the lattice, and concomitantly reduce both the landing frequency and run length of the soluble kinesin-1 isoform KIF5B at the single-molecule level, but not those of KIF5C. Notably, these effects are spatially heterogeneous along individual microtubules, indicating that microtubule lattice structure, and its mechanical response to tension, is not uniform along the filament. In cells also we found that single microtubules can accommodate distinct, non-overlapping patches of MAPs with competing preferences for expanded and compacted lattice. We also developed a minimal theoretical model rationalizing for the experimental observations. The model, based on a previously developed theoretical framework proposing the existence of highly cooperative conformational transitions of the MT lattice, where αβ-tubulin can adopt discrete and distinct conformations and dynamically isomerize between them (17), qualitatively and quantitatively reproduces our observations.

## Results

### Tensile forces expand the microtubule lattice

To determine whether tensile forces can structurally deform the microtubule lattice, we used a dual-laser trap in which biotinylated, GMPCPP- and paclitaxel-stabilized microtubules were attached at both ends to streptavidin-coated, laser-trapped polystyrene beads — a configuration referred to as a microtubule dumbbell (27, 29) (Fig. 1A; Movie S1). One of the traps was steerable independently via the use of an acousto-optic deflector, allowing control of the applied tensile force by adjusting the distance between the two traps. Microtubules were sparsely labelled with streptavidin-coated quantum dots (QDs) and imaged using high-contrast epifluorescence illumination to measure with nm-resolution accuracy force-induced deformations of the microtubule lattice via changes in the relative distance between the QDs (Materials and Methods). Control experiments using surface-immobilized QDs confirmed that super-resolution tracking localizes positions with a precision below 3 nm (Fig. S1; *SI Appendix*).

**Figure 1.**
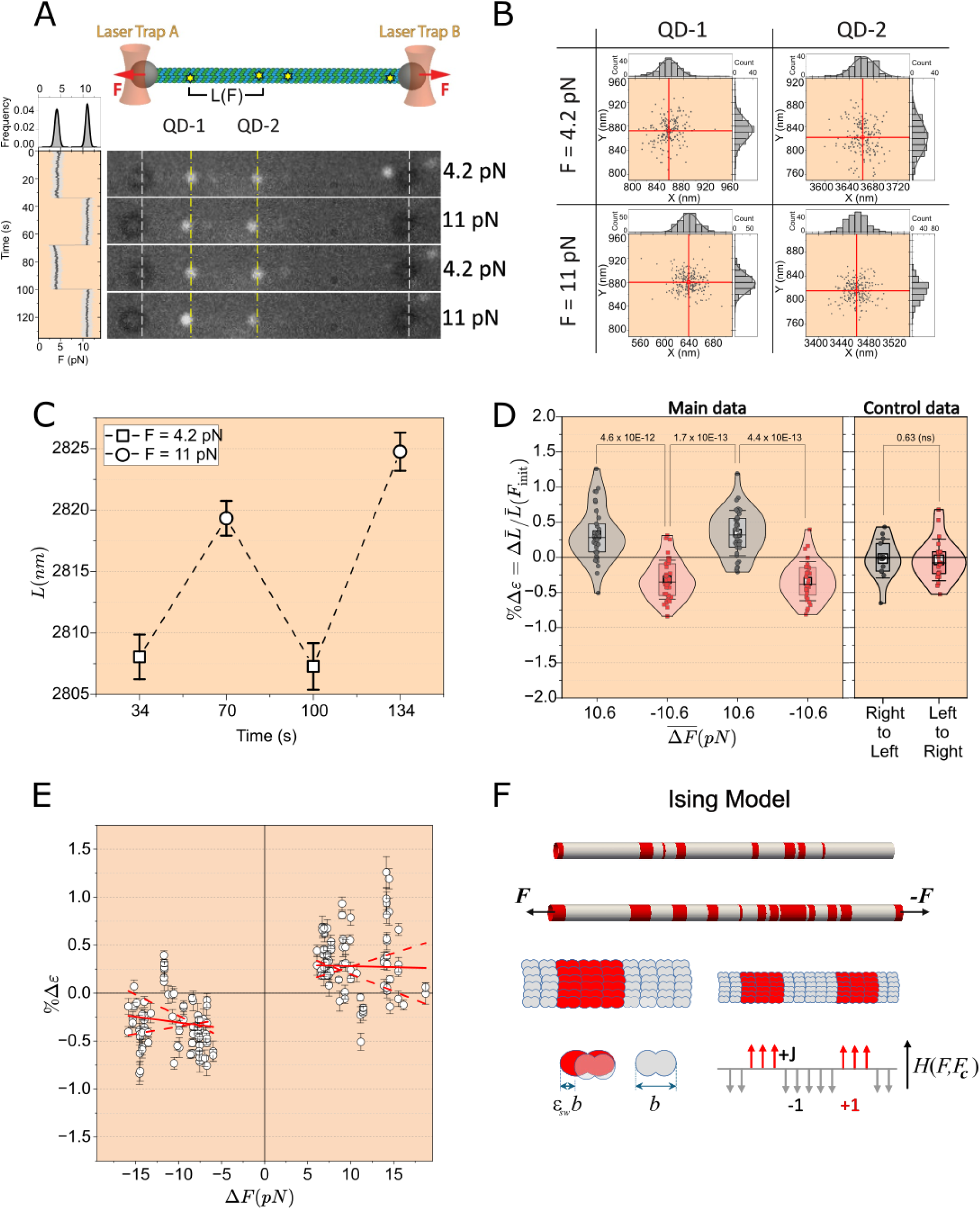
Microtubule lattice expansion under tensile forces is reversible and discrete, not elastic. (A) Top panel: Cartoon representation of a 50% biotinylated microtubule connected to two Neutravidin-coated polystyrene beads (gray spheres) at both ends. Yellow stars represent streptavidin-conjugated QDs attached to the microtubule. The two polystyrene beads are physically manipulated by two steerable optical traps. The magnitude of the tensile force, F, applied to the dumbbell is controlled by adjusting the position of trap A relative to trap B. The four panels below are single frames from 4 different 300-frame movies in which the dumbbell is subjected to different levels of tensile force indicated at the right of each. QDs are shown in bright color and the polystyrene beads in dark gray. Vertical dashed lines serve as a guide to the eye. On the left of the photos the force trace is shown as a function of time and on top of it a histogram of the force trace is shown. (B) Representative xy-localization distributions of QDs 1 and 2 shown in (A) at 4.2 pN and 11 pN. The vertical and horizontal red lines indicate the mean x- and y-position of each distribution, respectively. (C) The distance L between the mean positions of QDs 1 and 2 in (A) as a function of time and tensile force. (D) Distributions of the strain ε over four cycles of average change in tensile force 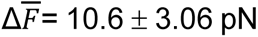 (SD). Statistical comparisons were performed using the two-tailed Mann-Whitney U test, with exact p-values reported. (Main Data:18 dumbbells 71 QD pairs; Control Data: N = 14 (gray), N = 21 (red)). (E) Scatter plot of the change in strain *Δε* as a function of the change in tensile force *ΔF* (N = 142 data points). The solid red lines for ΔF > 0 and ΔF < 0 represent linear fits calculated from the mean slope and corresponding intercept averaged over 10,000 bootstrap replicates, each median-filtered with a sliding window of 31 points. The dashed red lines correspond to the 2.5th and 97.5th percentiles of the bootstrap linear fit values, defining the 95% CI. Median filtering was applied before fitting to mitigate the disproportionate influence of regions with higher data point density on the slope estimation. (F) Schematic of the 1D Ising chain model, in which the microtubule lattice is represented as a series of effective sections shown in gray and red. Gray and red correspond to segments in the compacted (s = −1) and expanded (s = +1) conformational states, respectively. The length of a tubulin dimer in the expanded state is b(1+*ɛ*_*sw*_), where b is the dimer length in the compacted state and *ɛ*_*sw*_ is the switching strain. Application of a stretching force F biases the conformational equilibrium toward the expanded state, increasing the proportion of effective sections in the s = +1 state. H (F, F_c_) is the effective bias energy between the two conformational states, when F > F_c_ tubulin adopts the s = +1 and when F < F_c_ it adopts the s = -1 state (see text). Error bars in (C) and (E) were calculated using standard error–propagation formulas based on the super-resolution positional tracking uncertainty of each quantum dot.

Microtubule dumbbells decorated with QDs were subjected to cycles of increase and decrease in tensile force *F* in the range of *F*_min_ = 1.29 pN to *F*_max_ = 22.3 pN, and the distance between the mean positions of QD pairs was measured at each force level over ∼300 imaging frames (60 s). A visual example of a QD pair, their super-resolution position tracking, and their relative distance *L* as a function of tensile force applied to the microtubule are shown in Fig. 1A–C (see also Movie S1). Measurements were performed on 18 microtubule dumbbells and 71 QD pairs. For each change in force, the corresponding change in distance, *ΔL*, was calculated, and the resulting change in strain was approximated as Δ*ε* = Δ*L*(*F*)/*L*(*F*_init_) = [*L*(*F*_fin_)-*L*(*F*_init_)]/*L*(*F*_init_) for at least two cycles of change *ΔF* in force (Fig. 1D; for more details in the calculation of Δ*ε* see Supplementary Methods and Fig. S2). Dividing by the initial length *L*(*F*_init_) is a good approximation for the unloaded microtubule length *L*_0_ in the definition of strain ε = Δ*L*/*L*_0_ because strain values are in the sub-percent range. We plotted the change in strain values Δ*ε* for all 18 microtubule dumbbells as a function of the average increase and decrease of force <*ΔF*> = 10.6 ± 3.06 pN (SD) over four cycles (Fig. 1D left panel). The resulting chang in strain Δ*ε* is overall reversible and the distributions of its values during increase and decrease of the tensile force are statistically significant from each other (two-tailed Mann-Whitney U test, p<0.001). The average change in strain values for positive and negative change in force were (0.33 ± 0.080)% and –(0.33 ± 0.070)% respectively (errors represent 95% confidence intervals). As can be seen in Fig. 1D (left panel), not all Δε values are positive when *ΔF* > 0, nor are they all negative when *ΔF* < 0. We reasoned that this variability in the sign of Δε could reflect positional uncertainty due to (i) fluctuations in QD position relative to the microtubule lattice superimposed on the positional fluctuations of the microtubule dumbbell itself, and possibly (ii) an intrinsically fluctuating response of the microtubule lattice under constant force. To assess these contributions, we performed control experiments in which dumbbells were displaced laterally by ∼200–300 nm to the left or right by steering both traps simultaneously, while keeping the trap separation constant and therefore maintaining the same tensile force. These displacements mimic the lateral motions that accompany force increases and decreases in the main experiments (Fig. 1A). Under these conditions, the change in strain Δε did not differ significantly from zero in either direction, and its values remained within −0.5% to 0.5% (Fig. 1D, right panel), matching the range over which sign ambiguity in Δ*ε* is observed in the main dataset (Fig. 1D, left panel). In addition, control measurements of the relative distance changes *ΔL*/*L* between surface-immobilized Qdots during lateral motion of the surface via a piezo-stage showed an even narrower distribution of, within −0.2% to 0.2% (Fig. S3). Together, these controls indicate that the observed variability in the sign of Δε does not arise from imaging or illumination artifacts but instead reflects positional uncertainty. The possibility that changes in QD interdistances arise from force-induced global straightening of the microtubule was excluded, as the measured change in strain showed no dependence on the initial spacing between QD pairs (Fig. S5; *SI Appendix*).

Taken together, these results demonstrate that changes in tensile force *ΔF* as small as 10 pN, which would correspond to one or two kinesin motors, is sufficient to induce a longitudinal extension of the microtubule lattice, that is reversible on average, by ∼0.3%.

### Ising model

To quantify the mechanical properties of the microtubule lattice, the change in strain *Δε* was plotted as a function of the change in tensile force *ΔF* (Fig. 1E). Within the force range explored here (< 23 pN) and for a two- to three-fold change in force magnitude, Δ*ε* does not vary linearly with *ΔF* but instead displays a step-like transition between -(0.33 ± 0.35)% and (0.33 ± 0.28)% (errors are SD) depending on the sign of the force change *ΔF* (see fit in Fig. 1E). The slope of *Δε* vs. *ΔF*, calculated from 10000 bootstrap replicates, was not significantly different from zero for either *ΔF* < 0 (slope = -0.012 pN^-1^, 95% CI [-0.046, 0.15]) or *ΔF* > 0 (slope = -0.0019 pN^-1^, 95% CI [-0.035, 0.0029]), consistent with a step function response rather than a linear elastic response. Consistently, the intercepts of the linear fits were statistically significantly different from zero for both ΔF < 0 (*Δε*(*ΔF* = 0) = −0.43%, 95% CI [−0.73, −0.21]) and *ΔF* > 0 (*Δε*(*ΔF* = 0) = +0.30%, 95% CI [+0.066, +0.58]), confirming that the lattice adopts two distinct strain states rather than varying continuously around zero.. This implies a binary mechanical transition of the microtubule lattice, in which case the change in strain *Δε* and the strain *ε* itself are equivalent by definition, since |*ε*| can take only two values: 0 and 0.33%. Assuming the deformation is strictly elastic yields an anomalously soft modulus of ∼11 MPa (*SI Appendix*), which is more than two orders of magnitude below the expected Young’s modulus ∼1 GPa for the tubulin lattice (30). Interpreting it instead as a shear modulus leads to the equally unphysical conclusion that protofilaments along a 15 µm microtubule would slide past each other by ∼6 heterodimer lengths (*SI Appendix*).

The two-state nature of the microtubule strain response to applied force is consistent with a polymorphic lattice model in which tubulin heterodimers switch between two discrete conformational states, an expanded and a compacted one (17, 30). Since neighbouring tubulin heterodimers are not mechanically independent, we coarse-grain the cylindrical lattice into effective sections, each comprising a group of mechanically coupled heterodimers. Due to the cylindrical geometry of the microtubule and the applied uniaxial tensile force, all tubulin heterodimers within a given cross-section experience the same mechanical environment and are therefore assumed to adopt the same conformational state simultaneously. This reduces the problem to one spatial dimension: the effective sections have no azimuthal variability, their radius is fixed, and their length is the tubulin dimer size in the given conformational state. Each effective section thus occupies exclusively one of the two states at any given time. In the simplest limit of this 1D model (31), where high tension effectively suppresses the compacted state, we can model a microtubule under tension as an 1D Ising chain of such effective sections, each existing in one of only two discrete states *s* = +1, −1 corresponding to the expanded or long and compacted or short state, respectively (Fig. 1F). In this case (32) we write the conformational energy of a microtubule lattice under tensile force as the sum over all its effective sections *i*

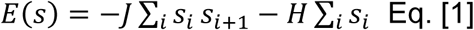

with the Ising ferromagnetic type coupling energy *J* > 0 favoring neighboring sections being in the same state (33) and 2*H* the effective energetic bias energy between the long and the short state. This bias energy *H* reflects the competition between the intrinsic free energy Δ*G* between the two conformational states in the absence of force and the mechanical work *F*×D*l* performed by the external tension *F* coupling to the elongation between the two states *Δl*;

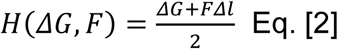

where Δ*l* = *bε_sw_*, with *ε_sw_* switching strain, and *b =* 8 nm the size of the tubulin heterodimer in the absence of force. Note that for the short state to be preferable in absence of force, *ΔG* < 0, while a sufficiently large tension *F* > 0 can flip the sign of the effective field *H*. The flipping happens when *F* exceeds the critical force *F*_c_ = *ΔG*/*Δl* for which *H*(*ΔG*, *F*_c_) = 0. The analogy with the Ising chain (*SI Appendix;* Table S1) gives rise to the force-strain relation

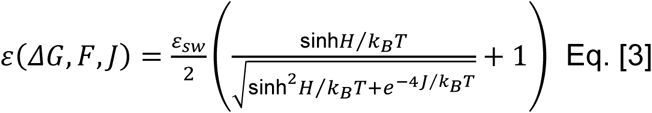

where *k_B_* is the Boltzmann constant and *T* the absolute temperature. Fitting the change in strain *Δε* as a function of *F*_initial_ and *F*_final_, i.e., *Δε* = *ε*(*ΔG, F_final_, J*) - *ε*(*ΔG, F_initial_, J*), using Eq. [3] (Fig. S6A) yields an average strain *ε_sw_* =(0.339 ± 0.056)% and a critical force *F_c_* = (8.44 ± 1.52) pN (95% CI), which correspond to an intrinsic free energy *ΔG* = 0.23 k_B_T. 5.9 *k_B_T*. The coupling energy J cannot be accurately determined from the data because, once the transition becomes sufficiently sharp — as is the case here — the predicted force-strain curve becomes indistinguishable from a perfect step function, rendering the data insensitive to the precise value of J. To determine a lower bound on J, we computed the fit residuals as a function of J while optimizing all other parameters (Fig. S6B). The residuals plateau at their minimum value for J ≳ 3 k_B_T, confirming that the data cannot distinguish between values of J above this threshold. The residuals increase monotonically for J < 3 k_B_T, establishing J = 3 k_B_T as a lower bound on the coupling energy. For J < 0 the residuals reach a second plateau at substantially higher values, confirming that J > 0 is strongly favoured by the data. The robust determination of *F*_c_ and *ε*_sw_ confirms that the data quality and fitting procedure are sound. It should also be noted that these represent ensemble-level values, and the parameters extracted for individual microtubules can be considerably variable, as reflected by the scatter of the data around the mean. The standard deviation of the strain is 0.35% for positive values and 0.28% for negative values.

### Tensile forces on the microtubule lattice affect its interaction with kinesin1

If the force-induced longitudinal expansion of the microtubule lattice reflects structural changes in its architecture, it should modulate the biochemical interactions of MAPs or motor proteins, such as kinesin1. Previous work by some of the authors showed that when both kinesin and microtubule are under forces that oppose their relative motion not all areas of the same microtubule are equivalent interacting substrates for truncated human kinesin1 isoform KIF5B (hKIF5B(560)) commonly known as K560 (27). The median attachment duration of kinesin to the microtubule can differ by more than an order of magnitude within different locations on the same microtubule or between different microtubules and switch between catch- and slip-bond type of behaviour. This effect however is absent for soluble kinesin1 on force-free microtubules.

To test whether the interaction of soluble fluorescently tagged hKIF5B(K560)-mNeonGreen (hKIF5B(560)-mNG; see SI) at the single-molecule level is influenced by tensile forces applied to the microtubule lattice, we developed an assay using surface-tethered microtubules polymerized with GMPCPP from surface-immobilized GMPCPP seeds (see Materials and Methods). The microtubules were subsequently capped at both ends with 10% biotinylated tubulin and further stabilized with paclitaxel. Tensile forces were applied by attaching a laser-trapped, streptavidin-coated polystyrene bead to either of the biotinylated ends (Fig. 2A; Movie S2). Combining optical trapping with highly inclined and laminated optical sheet illumination (HILO) (34) enabled simultaneous visualization of soluble single hKIF5B(560)-mNG molecules interacting with microtubules subjected to tensile force oscillated in a step-function fashion between zero and a non-zero value with a period of 120 seconds.

**Figure 2:**
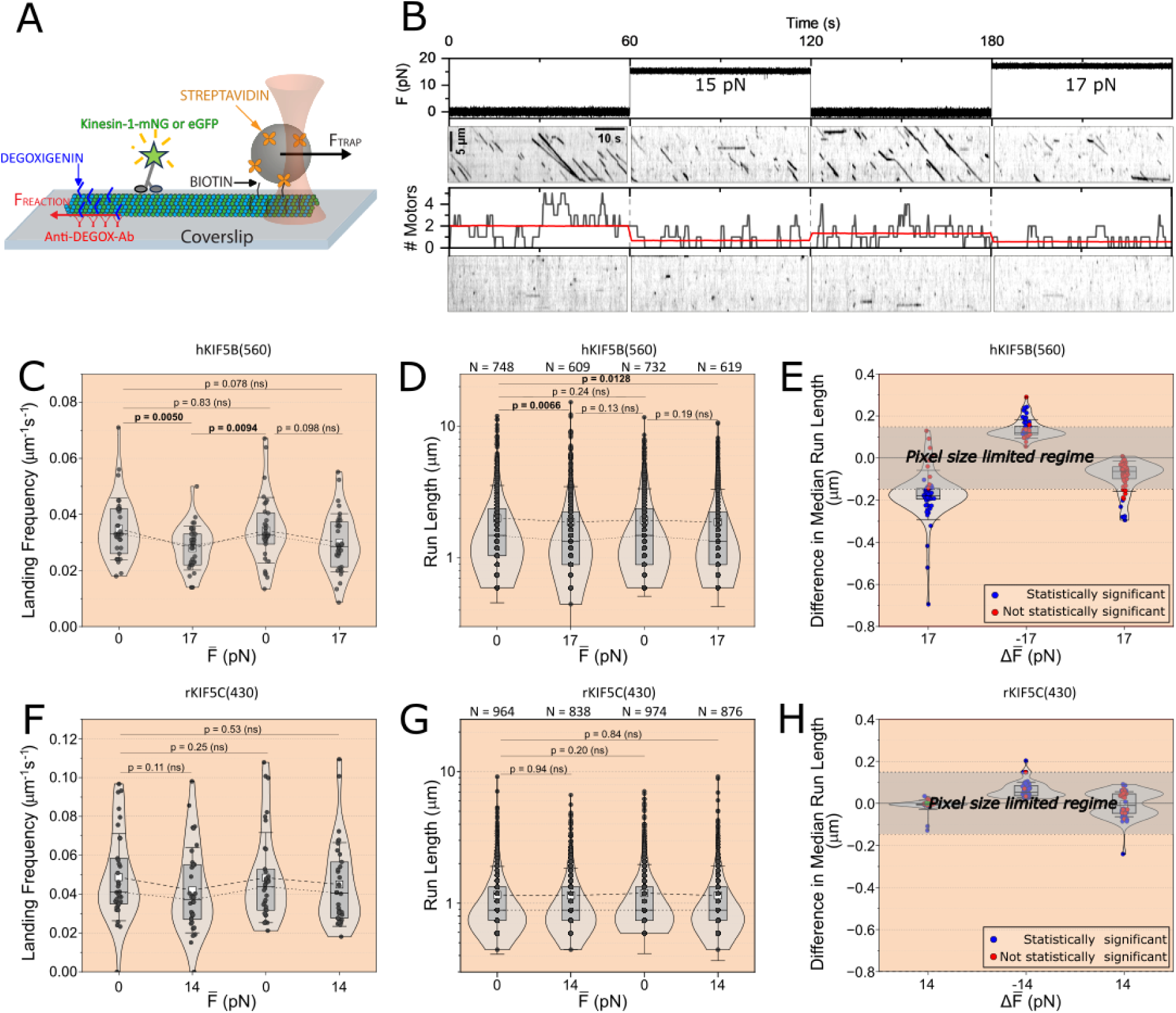
Expansion of the microtubule lattice affects its interaction with KIF5B but not with KIF5C. (A) Cartoon representation of the assay. A surface-tethered microtubule, anchored at its digoxigenin-labeled (– or +) end, is pulled at its biotinylated (+ or -) end via a laser-trapped streptavidin-coated bead. Coupling with HILO illumination enables single-molecule visualization of hKIF5B(560)-mNG while the microtubule is under tensile load. (B) Top: Representative force trace over time. Upper Middle: Kymographs of hKIF5B(560)-mNG runs corresponding to the indicated force regimes. Lower Middle: Number of running motors along the microtubule as a function of time (black line). The red line shows the mean number of motors. Bottom: Kymograph of background fluorescence recorded along a line parallel to, and 10 µm away from, the microtubule under study. (C) Violin–box plots of the landing frequency of hKIF5B(560)-mNG where each point is a microtubule. A statistically significant ∼20% decrease (exact p-values; ns stands for non-significant; two-tailed Mann-Whitney U test) in the median landing frequency is observed under an average tensile force of 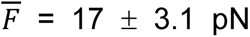 (28 microtubules; F_max_ = 25.2 pN, F_min_ = 9.71 pN). P-values lower than 0.05 are indicated in bold. The dashed and dotted lines connect the average and median values of each distribution, respectively. (D) Violin–box plots of the corresponding run lengths show a statistically significant ∼10% decrease (asymptotic p-values, due to large sample size; two-tailed Mann-Whitney U test) in median run length during the first force cycle under the same average force. The number of single-molecule tracks for each force is indicated at the top of the plot. The discrete appearance of the scatter plot reflects the pixel-limited resolution of the tracking data, as run lengths are integer multiples of the camera pixel size. Run length distributions appear non-exponential due to truncation of tracks at the boundaries of the microtubule segment between the seed and the cap (see *Materials and Methods*) (E) The *difference of the median run length* upon the average *change of force* 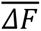 was calculated for various microtubule subsets by sequentially excluding microtubules whose removal maximizes or minimizes the difference in the median run length. Statistically significant differences (95% CI) are shown in blue and non-significant differences are in red. The resolution limited regime due to the pixel size of the camera (148 nm) is indicated by the gray transparent band. (F)-(G) panels are the corresponding panels to (C)-(E) but for rKIF5C(430)-eGFP. The errors in all the box plots correspond to the standard deviation, the horizontal line to the median, the open square to the mean and the box width to the central 50^th^ percentile of each distribution.

The lowest applied force was always between 0 pN and 1 pN, while the maximum force ranged between 10 pN and 25 pN. The minimum non-zero tensile force was set to 10 pN, since our previous QD-based measurements detected microtubule deformations when force is higher than F_c_ = 8.4 pN. Because kinesin-1 motors have been shown to communicate through the microtubule lattice (35, 36), experiments were performed at sub-nanomolar motor concentrations to allow single-molecule observations and minimize potential microtubule lattice-mediated crosstalk. A representative force trace and the corresponding kymographs for hKIF5B(K560)-mNG are shown in Fig. 2B.

Quantification from 28 individual microtubules revealed that applying tensile load resulted in a statistically significant reduction (two-tailed Mann-Whitney U test) in both the median landing rate (s⁻¹·µm⁻¹) and the median run length of hKIF5B(560)-mNG by ∼20% and ∼10%, respectively (Fig. 2C, D). Upon increasing the average tensile force to 17 pN, the reductions in median landing rate and median run length, and their subsequent recovery upon returning to 0 pN, did not remain statistically significant over two successive force cycles, suggesting that the mechanical response of the microtubule lattice attenuates upon repeated force cycling.

However, substantial microtubule-to-microtubule variability was observed, with not all microtubules responding similarly to the application of tensile force. This heterogeneity does not depend on the magnitude of the tensile force (Fig. S7) and is better illustrated by calculating the median run length for different subsets of the data through a serial exclusion analysis, in which microtubules were removed one at a time from the full dataset (N = 28) in a manner that maximized (or minimized) the median run length of the remaining *N − i* microtubules, where *i* denotes the iteration number of serial exclusion. Then for each subset the changes in median run length over successive cycles of changing force were calculated (Fig. 2E). As can been seen the magnitude of the effect of tensile forces on hKIF5B(560)-mNG’s run length is highly variable depending on the subset of microtubules. Interestingly this heterogeneity does not exhibit any statistically significant correlation with the magnitude of the average tensile force of each subset (Fig. S8). The diminishing response in the change of median run length can again be observed over successive cycles of force application. No statistically significant change in hKIF5B(560)-mNG velocity was observed under tension (Fig. S9).

Analogous experiments were performed with the fluorescent truncated rat kinesin-1 isoform KIF1C-eGFP (rKIF5C(430)-eGFP), known to expand GDP-lattices when bound in rigor (absence of ATP) (20, 21). In contrast to hKIF5B(560)-mNG, rKIF5C(430)-eGFP showed overall no statistically significant changes in either landing rate or run length across the similar tensile-force range (Fig. 2F-G). Statistically significant changes in run length were observed only in rare instances within a small subset of the data, and when present, they occurred in the same direction as observed for hKIF5B(560)-mNG, i.e., a decrease in run length upon application of tensile force (Fig. 2H).

Together, these results indicate that mechanical tension on microtubules can regulate its interaction with kinesin-1 in an isoform-dependent manner and that this response can vary across individual microtubules *in vitro*.

### The microtubule lattice is a dynamic mosaic of conformational states

We hypothesised that the observed heterogeneity in mechanical response across microtubules (Fig. 2D, E) reflects differences in the conformational states of the microtubule lattice. Since mechanical forces can alter the lattice, we asked whether repeated cycles of tensile force application might shift the lattice into new conformational states, generating heterogeneity over time within individual microtubules. To explore this, rather than measuring the median run length, we quantified the running motor density averaged over the time of the kymograph, <ρ>_t_ (µm⁻¹) of kinesin molecules along individual microtubules over multiple force cycles. This metric accounts for both the landing frequency and dissociation rate of kinesin along different microtubule segments, and describes how frequently any given segment is permissive or non-permissive to hKIF5B(560)-mNG landing and processive movement. Due to the low kinesin concentration, the number of events per kymograph (1 min) can fall below 30. To enable statistical comparisons across different force values, running motor density profiles were therefore calculated by pooling two consecutive kymographs (2 min) at each force level.

The running motor density profiles for the microtubule shown in Fig. 2B are presented in Fig. 3, together with the corresponding kymographs, for zero and non-zero force over four cycles of force application. During the first two cycles (t < 240 s), the average motor density at zero force (0.13 µm⁻¹) is significantly higher (95% CI) than that measured at an average force of 16 pN (0.051 µm⁻¹) (Fig. 3B). At both zero force and 16 pN, the motor density is uniform along the microtubule, since the variances of the corresponding motor density profiles are not significantly different from that expected for a uniform distribution of motors along the microtubule (95% CI) (see *Supporting Information*: Analysis of fluorescent movies, kymograph quantification and running motor density profile). In the subsequent two pulling cycles (t > 240 s) (Fig. 3C), however, the motor density profiles deviate significantly from a uniform distribution (95% CI). At 17 pN, the minus-end proximal segment (L < 7 µm) shows significant hKIF5B(560)-mNG enrichment relative to the plus-end proximal segment (L > 7 µm). Furthermore, the difference in average running motor density remains almost the same at zero (0.092 µm^-1^) and non-zero force (0.082 µm^-1^). To exclude the possibility that repeated pulling caused lattice damage, control experiments were performed in the presence of 2% fluorescently labelled tubulin (Alexa 488). No tubulin incorporation into the lattice was detected over at least 6 pulling cycles (Fig. S10). Together, these observations indicate that the conformational state of the microtubule lattice depends on its loading history. Although force returns to zero after each pulling cycle, the lattice does not necessarily recover its original conformation within the 1 min half-period timeframe.

**Figure 3:**
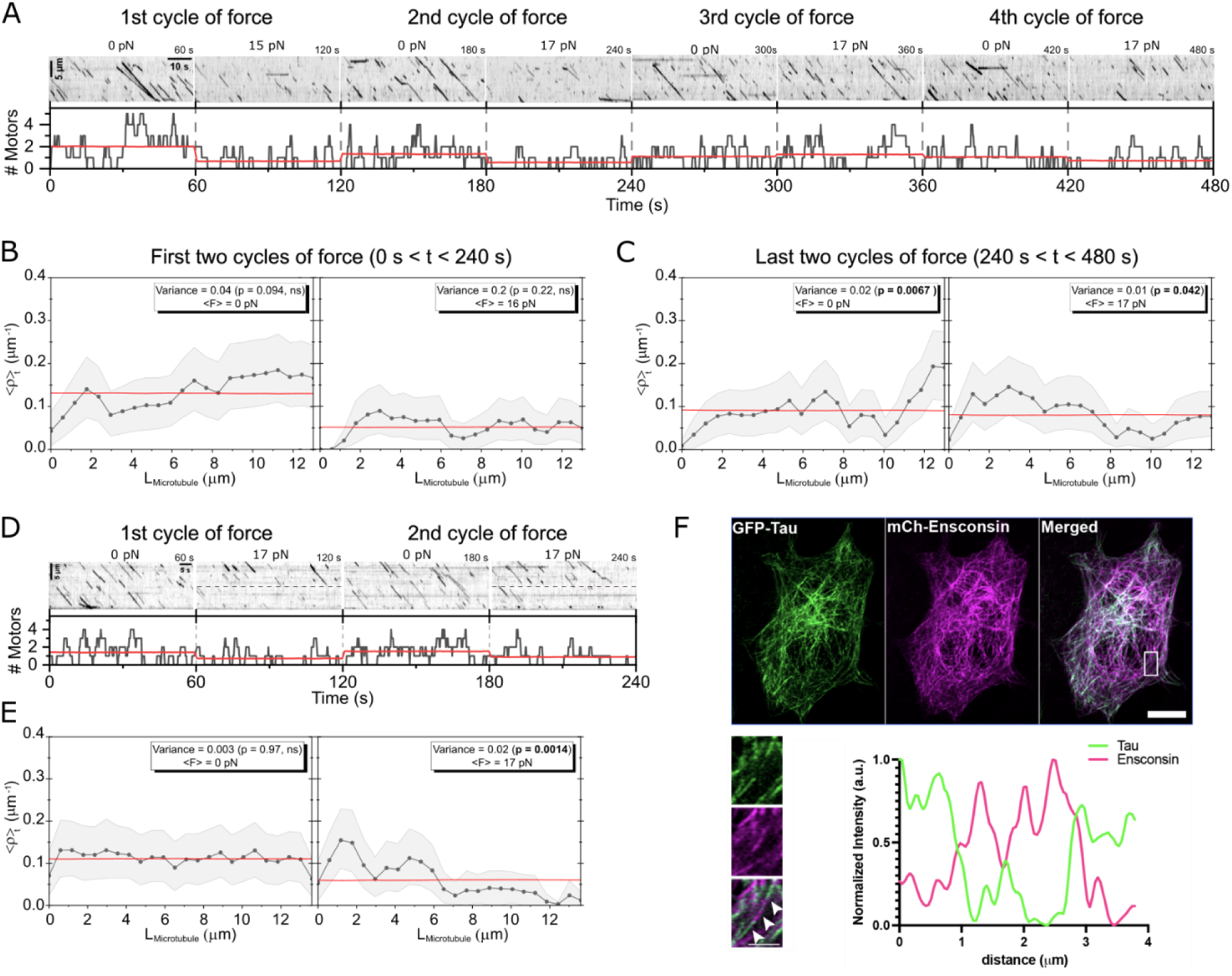
Heterogeneity along individual microtubules *in vitro* and in cells. Kymographs of the same microtubule as in Fig. 2B over four force cycles in which the force level is changed every minute. Below, the corresponding number of running motors as a function of time is shown (black line). The red line represents the expected number of motors assuming a uniform distribution over time (see Fig. 2B). (B) Time-averaged running motor density profiles <ρ>_t_ for the first two force cycles (t < 240 s). Due to the low hKIF5B(560) concentration and the consequently small number of events per kymograph, two consecutive one-minute kymographs were pooled at each force level to enable statistically meaningful comparisons (1st and 3rd minutes at ⟨F⟩ = 0 pN; 2nd and 4th minutes at ⟨F⟩ = 16 pN). The black scatter line shows the <ρ>_t_ (µm⁻¹) as a function of distance from the microtubule minus end. The shaded band indicates the 95% CI calculated from 10^4^ bootstrap replicates. The red line represents the time-averaged motor density obtained from 10^4^ bootstrap replicates generated under the null hypothesis that all positions along the microtubule are equally accessible for motor landing and processive runs. The variance of the <ρ>_t_ profile is compared to that of the bootstrap null distribution, and statistical significance is assessed using the 95% confidence interval. P-values lower than 0.05 are indicated in bold. (C) <ρ>_t_ profiles for the same microtubule during the last two force cycles from (A). (D) Kymographs of a different microtubule subjected to alternating force levels of F = 0 pN and F = 17 pN applied every minute. The dashed line overlaid on the F = 17 pN kymographs serves as a guide to the eye, indicating the boundary beyond which hKIF5B(560) motor landing and processive runs are markedly reduced. (E) <ρ>_t_ profiles for the kymographs shown in (D). Statistical comparisons were performed as in (B). (F) Upper panel: Representative fluorescence images of the same HeLaM cell expressing eGFP-tau and mCh-ensconsin, shown individually and as a merged overlay (scale bar 10 µm). Lower panel: (left) Magnified view of the boxed region indicated in the upper panels (scale bar 1 µm). The three white arrows mark the microtubule successive segments along which the normalised intensity profile shown on the right was measured.

A similar spatial bias in hKIF5B(560) enrichment is shown for a second microtubule (Fig. 3D, E). Here, the hKIF5B(560)-mNG density profile is uniform at zero force but statistically significantly enriched in the minus-end proximal segment (L < 6 µm) relative to the plus-end proximal segment (L > 6 µm) at a pulling force of 17 pN.

We next asked whether the structural heterogeneity observed along single microtubules *in vitro* is also present in cells. To address this, we co-expressed fluorescently tagged MAPs, mCherry–ensconsin (MAP7) and eGFP-tagged full-length 2N4R tau, in HeLaM cells. Tau has been associated with compacted microtubule lattice states, whereas ensconsin has been proposed to preferentially associate with expanded or stable lattice conformations (37, 38). As shown in Figure 3F (panel “Merged”), tau-positive filaments did not overlap with ensconsin-labeled microtubules, indicating spatial segregation between MAPs with distinct lattice preferences. Upon closer inspection, a subset of microtubules exhibited both tau and ensconsin localization along the same filament (Figure 3F), however, these proteins occupied distinct, non-overlapping regions, as confirmed by line intensity analysis (Figure 3F, graph). This alternating pattern suggests that individual microtubules can contain spatially distinct lattice states that differentially recruit MAPs. Importantly, treatment with 10 µM paclitaxel, which is known to expand microtubules (39, 40), disrupted linear tau-decorated filaments (Fig. S11), with tau redistributing into a diffuse cytoplasmic pool, whereas ensconsin-labeled microtubules remained intact.

Together, these data support the view that individual microtubules, both *in vitro* and in cells, are not always structurally uniform. Instead, they comprise a mosaic of regions with distinct conformational states that can be dynamically modulated by tensile forces or MAPs.

### Simulations of the microtubule lattice dynamics

To explain the experimental observations of the dynamic kinesin force response within the framework of the proposed two-state polymorphic lattice model, we performed Glauber-Ising-type dynamic simulations in which the lattice state responds dynamically to applied tension, coupled to agent-based kinetics of kinesin motor binding, unbinding and stepping along the switchable lattice (Fig. 4A–D). The applied tensile force was oscillated in a step-function fashion between 0 and 17 pN with a period of 120 s, while the critical force F_c_ was set to 8.4 pN as determined from the fit to the strain data (Fig. S5). Kinesin motors are assumed to bind and step toward the microtubule plus end exclusively when in contact with the compacted lattice state (white state, s = −1). Upon encountering an island of expanded-state tubulin (red state, s = +1), motors spontaneously unbind and return to solution (Fig. 1F). When the applied force exceeds the critical conversion force (F > F_c_), the lattice, initially residing in the compacted, high-affinity state for kinesin, progressively converts to the expanded, low-affinity state (Fig. 4A). This progressive transition is quantified by the spatially averaged tubulin state ⟨s⟩, which ranges from −1 (all tubulins compacted) to +1 (all tubulins expanded) as a function of force and time (Fig. 4B). The corresponding profiles of the number of motors over time and averaged motor density ⟨ρ⟩_t_ along the microtubule are shown in Figs. 4C-E.

**Figure 4:**
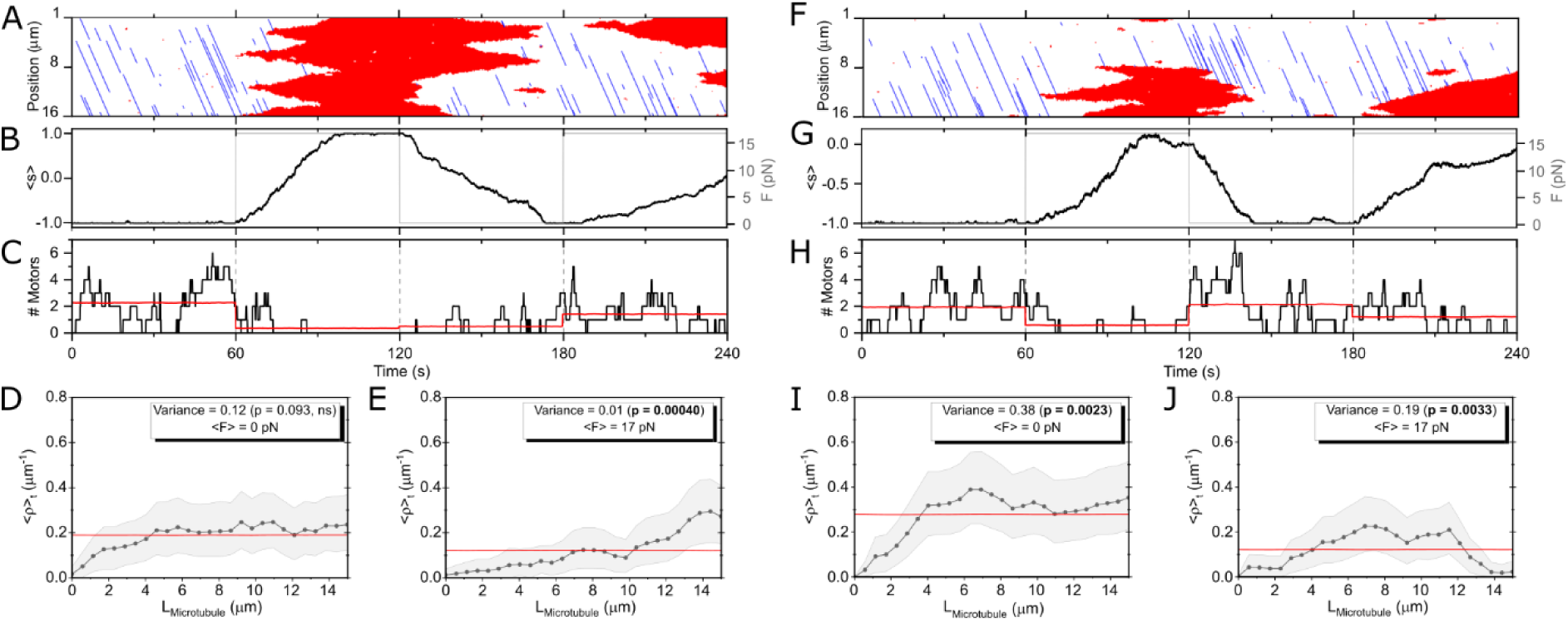
Kymographs and time-averaged motor density profiles from Glauber simulation using the 1D Ising model. (A–E) Simulation of lattice state and kinesin dynamics for a homogeneous microtubule lattice of length L = 16 µm with a uniform critical force F_c_ = 8.5 pN (corresponding to a bias field H = −0.03 *k_B_T*). (A) Time evolution of kinesin positions (blue lines) and microtubule lattice states (red areas) for the homogeneous lattice. (B) The applied force alternates between 0 pN and 17 pN over two complete cycles of 120 s each (total duration 240 s; grey line, right y-axis). The mean tubulin conformational state ⟨s⟩ (black line, left y-axis) responds by switching between the compacted (⟨s⟩ = −1) and expanded (⟨s⟩ = +1) states. (C) Time evolution of the total number of microtubule-bound kinesin motors. (D) Time-averaged running motor density profile ⟨ρ⟩_t_ (grey line and points) for the two force-free sub-cycles (F = 0 pN), which is not statistically different from a uniform distribution (red line; 95% CI). (E) ⟨ρ⟩_t_ profile for the two high-force sub-cycles (F = 17 pN), which deviates significantly from a uniform distribution (red line; 95% CI). (F–J) Simulation of lattice state and kinesin dynamics for a heterogeneous microtubule lattice of length L = 16 µm comprising two segments of equal length — a minus-end proximal segment (0–8 µm) with high critical force F_c1_ = 20.4 pN and a plus-end proximal segment (8–16 µm) with low critical force F_c2_ = 8.5 pN. (F) Time evolution of kinesin positions (blue lines) and microtubule lattice states (red areas) for the heterogeneous two-block lattice. (G) Time course of applied force and mean lattice state ⟨s⟩ for the heterogeneous lattice (corresponding to panel F). Note that ⟨s⟩ does not reach +1 upon force application because the plus-end proximal segment (8–16 µm), with F_c2_ = 20.4 pN, remains in the compacted state throughout, limiting the maximum mean lattice state to values close to 0. (H) Time evolution of the total number of bound kinesin motors (corresponding to panel F). (I) ⟨ρ⟩_t_ profile for the two force-free sub-cycles (F = 0 pN), which deviates significantly from a uniform distribution (95% CI). (J) ⟨ρ⟩_t_ profile for the two high-force sub-cycles (F = 17 pN), which also deviates significantly from a uniform distribution (95% CI). The reduction in mean running motor density from 0.28 µm⁻¹ to 0.12 µm⁻¹ upon force application (panels I and J) is statistically significant (95% CI). Simulation parameters: Microtubule lattice: J = 3 *k_B_T*, *H* = −0.03 *k_B_T*; Glauber-Ising update rate: 5000 steps s⁻¹. Kinesin motors: 50 motors dynamically exchanging between microtubule-bound and unbound states with binding rate p_bind_ = 10⁻¹ s⁻¹, corresponding to a landing rate of 0.01 µm⁻¹ s⁻¹ when normalized to the microtubule length (consistent with Fig. 2C and Fig. 2F), unbinding rate p_unbind_ = 1 s⁻¹ and stepping velocity v_run_ = 1 µm s⁻¹ (Fig. S8). The chosen unbinding rate yields an average run length of 1 µm. All 95% CIs were estimated from 10000 bootstrap replicates as in Fig. 3.

To account for the heterogeneous mechanical response along individual microtubule lattices observed in Fig. 2C–E, we modelled the microtubule as consisting of two segments of equal length but distinct critical forces, F_c1_ = 20.4 pN and F_c2_ = 8.5 pN, such that F_c1_ < F < F_c2_ (Fig. 4F–J). In both the uniform and heterogeneous simulations, application of force reproduces the experimentally observed suppression of motor binding and processive runs, as well as the statistically significant reduction in average running motor density (95% CI) and the statistically significant spatial bias in the time-averaged motor density profile ⟨ρ⟩_t_ (95% CI) (Fig. 4).

It is important to note that kinesin binding stabilizes the compacted tubulin state, while motors running into islands of expanded-state tubulin may actively perturb the local conformational dynamics of those islands, thereby influencing the overall evolution of the lattice. As a result, the observed mechanochemical response is not fully independent of the probe used to measure it: kinesin simultaneously reports on and modulates the conformational state of the lattice. This probe-dependent feedback is an inherent feature of the motor–lattice interaction rather than an experimental artefact, and is explicitly accounted for in the simulations, where kinesin kinetics and lattice dynamics are fully coupled.

## Discussion

Our findings establish that physiologically relevant tensile forces, in the range generated by one to three kinesin motors, can measurably and reversibly expand the microtubule lattice, resulting in modulation of the interaction between kinesin-1 and the microtubule at the single-molecule level. Notably, increases in tension < 20 pN reduce both the landing frequency and run length of the truncated (1-560) human kinesin-1 isoform KIF5B-mNG in a mostly reversible manner, whereas truncated (1-430) rat kinesin-1 isoform KIF5C-eGFP is largely insensitive under the same conditions, indicating isoform-specific mechanosensitivity of kinesin–lattice interactions. Moreover, this modulation is spatially heterogeneous along individual microtubules, implying that distinct regions can exist within the same microtubule that respond differently to mechanical tension and therefore do not constitute equivalent interaction substrates for kinesin-1.

These observations suggest that tubulin, even at a fixed nucleotide state (here GMPCPP), can adopt a more expanded conformation within the microtubule lattice when subjected to mechanical tension. Indeed, for each nucleotide-analogue state — GMPCPP, GTPγS, and GDP — the microtubule lattice has been shown to span a range of lattice spacings depending on whether, and which, MAPs or motor proteins decorate it (6, 23). The ∼0.3% expansion observed here for GMPCPP microtubules under an average tensile force increase of ∼10 pN falls well within the ∼1.4% range of lattice-spacing variation reported across available GMPCPP structures decorated with different MAPs and kinesin-1 motors, or left undecorated (6, 23). Furthermore, the decrease in both landing frequency and run length of hKIFB(560)-mNG upon force-induced lattice expansion is consistent with mechanically induced structural changes at the level of the αβ-tubulin dimer. Notably, hKIF5B(560) has been shown to compact the GMPCPP lattice by ∼0.79 Å (∼0.9%) (6); tensile forces would therefore be expected not only to oppose this compaction but also to bias the lattice toward a more expanded state, potentially shifting it away from the spacing optimal for hKIFB(560)-mNG binding and processivity. The observed reduction in landing frequency and run length of hKIFB(560)-mNG on GMPCPP microtubules under tension is thus consistent with this picture.

By contrast, no strong force-dependent effect was observed for rKIF5C(430)-eGFP (rat kinesin-1, isoform KIF5C) in the present study. This may indicate that rKIF5C(430)-eGFP is sensitive only to forces exceeding ∼20 pN (corresponding to greater lattice expansion) and/or that any force-induced reduction in run length falls below the spatial resolution limit of the present measurements (see Materials and Methods). It is noteworthy that, although rKIF5C(430) has been shown to expand the GDP microtubule lattice toward a GMPCPP-like conformation when bound in the apo state (20, 41), cryo-EM studies using human hKIF5B(560) report minimal structural changes in the GDP lattice upon kinesin binding (6). Together, these observations suggest that mechanosensitivity of the microtubule lattice within the kinesin-1 family may be isoform-specific.

The most surprising finding of our study is the heterogeneity of hKIFB(560)-mNG interaction in response to mechanical tension, observed both across different microtubules and along the length of individual ones, an observation that implies conformational multiplicity of the microtubule lattice. Closer inspection of the data reveals that distinct regions along a single microtubule can respond differently to applied tension, as evidenced by statistically significant spatial variations in kinesin hKIFB(560)-mNG time-averaged motor density profiles <ρ>_t_ along the microtubules (Fig. 3). Furthermore, upon successive cycles of force application and release, individual microtubules appear to adopt different conformational states between later timepoints at the same force level, as reflected in changes of <ρ>_t_ profile along the lattice. Because our *in vitro* assays were performed using purified components, biochemical alterations of the lattice can be excluded, making conformational change the most parsimonious explanation for the observed variations in motor density profiles. Additionally, we showed that the coexistence of non-overlapping tau- and ensconsin/MAP7-positive regions along the same microtubule suggests that local variations in lattice spacing may exist within individual microtubules in cells. The mixed population was abolished upon paclitaxel addition, which displaced the compactor protein (tau) from the lattice into the cytosol while leaving the expander (ensconsin) unaffected. These spatially distinct lattice states could provide a mechanism through which local mechanical forces regulate MAP and motor recruitment along individual microtubules in a spatially restricted manner. This is consistent with recent work demonstrating that competition between paclitaxel, which stabilises the expanded lattice state, and doublecortin, which favours the compacted state, produces microtubules with partial and non-uniform doublecortin decoration in cells, an effect that depends on the ratio of paclitaxel to doublecortin concentrations (42).

What is the physical basis for such inter- and intra-microtubule heterogeneity? Several non-mutually exclusive mechanisms may contribute both *in vitro* and in cells. First, since brain tubulin was used in our *in vitro* assays, which contains a mixture of different isotypes, the formation of segments with distinct isotype compositions along individual microtubules, as recently demonstrated *in vitro* (43), cannot be excluded. Such segments may differ in their intrinsic mechanical properties or structural conformation and thus respond differently to applied tension. Second, even when hydrolysis-deficient mutants of pure recombinant human tubulin isotypes were used, GTP-state microtubule lattices were found to consist of a mixture of regions adopting at least two distinct conformational states, characterized by high or low affinity for microtubule end-binding protein 3 (EB3), with occasional spontaneous switching between them (23). This indicates that structural heterogeneity can emerge even from compositionally uniform tubulin. Third, it is well established that microtubules reconstituted *in vitro* as well as in cytoplasmic extracts harbor various structural lattice defects. This includes switches in protofilament number, variable helical start numbers, and variable number and location of seams, that give rise to discontinuities and structural heterogeneity along microtubules (44–48). Importantly, structural defects are not confined to *in vitro* preparations but have also been identified in microtubules in cells (49, 50). Although the impact of these defects in cells is not fully understood, they are increasingly recognized as functionally important for the passive or active recruitment of specific MAPs (51–57). In this context, heterogeneous response to mechanical tension is most probably a manifestation of the underlying structural heterogeneity of the microtubule lattice.

Drawing on the results of the current work, we developed a minimal statistical mechanical 1D model built on the concept of a conformationally bistable tubulin heterodimer, as previously proposed by Mohrbach et al. (58). In this framework, every tubulin αβ subunit, regardless of isoform identity or nucleotide state, can switch between a compacted and an expanded conformation relative to its neighbors. The precise values of these two conformational states may vary depending on the tubulin isoform and/or nucleotide state, and may in turn influence binding proteins (including kinesin motors) in a context-dependent and potentially subtle manner. Fitting our data for the paclitaxel-GMPCPP microtubule lattices yields a measurable change of lattice elongation by 0.34%, a value that is comparable in magnitude with the previous estimates for paclitaxel-stabilized GDP microtubule loops and rings in gliding assays (59). The finite coupling energy *J* (on the order of a few *k*_B_*T*) implies a conformational correlation length *ξ* ∼ exp(2*J/k*_B_*T*) (in units of heterodimer length; see Table S1) in the micron range. This is consistent with studies of surface grafted paclitaxel-stabilized GDP microtubules exhibiting anomalous cooperative shape fluctuations and forming detectable super-helices (30, 60), but also with the long-range microtubule mediated interaction of kinesin motors over several microns (35, 36). Perhaps the most surprising, yet experimentally robust, finding is the small critical force of 8.5 pN required to induce the discrete lattice transition. Associated with this is a remarkably small free energy difference between the two conformations of ∼0.2 k_B_T. Notably, this critical force is smaller than the ∼ 30 pN required to extract a tubulin dimer from the microtubule lattice when pulling by its α-C-terminal tail (61). It nevertheless falls well within the range of stretching forces (2–15 pN) known to induce conformational changes in other mechanosensory proteins, such as talin and αE-catenin, which facilitate vinculin binding to reinforce focal adhesions and cell-cell junctions, respectively (62–65). The co-existence of two nearly isoenergetic conformational states — with free energy differences below k_B_T— is not without precedent. NMR has revealed that the R3 domain of talin co-exists in two distinct conformations in the absence of force (66), and single-molecule FRET has shown that the chaperon GroEL populates four distinct microstates with occasionally similar occupancies (ΔG << k_B_T), modulated by nucleotide state (67). In this context it is entirely plausible that fine-tuning of structural and energetic parameters allows tubulin to transition between two microstates upon very modest pulling forces equivalent to those generated by one or two kinesin motors. It is also worth noting that forces in the low pN range applied to filamentous actin, another major cytoskeletal filament, have been shown to regulate its interaction with soluble α-catenin (68).

We previously observed similar heterogeneity in which hKIF5B(560) attachment duration varied by more than an order of magnitude under opposing load at different locations along individual microtubule dumbbells (three-bead assay) (27). At the time, this striking variability — ranging from slip- to catch-bond type behaviour — was attributed to differences in shear stress arising from the fact that the two opposing forces, applied by surface-immobilised kinesin (F_kinesin_) and by the laser-trapped bead at the microtubule plus-end (F_trap_), could act along the same protofilament or along protofilaments separated by varying numbers of subunits around the microtubule circumference, generating different degrees of shear deformation of the lattice. In the present assay, however, the force pair — applied by the laser-trapped bead at the microtubule end (F_trap_) and by the surface-attached seed (F_reaction_) — acts on approximately diametrically opposite sides of the microtubule cross-section, separated by roughly half the circumference (Fig. 2A). Since this geometry is fixed and preserved across successive force cycles regardless of conformational changes, differences in the relative positioning of the two forces around the microtubule circumference can be excluded as the source of the heterogeneous KIF5B(560) response observed both here and in our earlier work. Furthermore, post-translational modifications (PTMs) of the tubulin C-terminal tails do not appear to contribute to this heterogeneity. Following subtilisin cleavage of the tails, hKIF5B(560) still exhibits both slip- and catch-bond behaviour at different locations along the same microtubule lattice in the three-bead assay (Fig. S12). Note that this treatment cannot be applied in the assay of Fig. 2A as subtilisin would indiscriminately cleave the anchoring antibodies and blocking reagents as well. In light of the above, the most parsimonious explanation for this variability is the dynamic polymorphic nature of the microtubule lattice and its modulation by tensile force.

There are many instances in cells where microtubules are known or have been proposed to be under tension. During mitosis, astral and spindle microtubules are actively pushed and pulled by motor proteins to position the spindle and define the division plane (69, 70). Dynein has also been implicated in placing microtubules under tension during microtubule organizing centre (MTOC) positioning in cells (71). Physical coupling of microtubules to the actomyosin network, for example at focal adhesions in migrating cells (72) or during axonal elongation in neurons (73), can potentially subject microtubules to tensile forces, although direct evidence for this remains to be established. Recently actomyosin contractility was found to drive submicron oscillations of microtubules along the axons of *Caenorhabditis elegans* neurons (74). Interestingly, microtubules have emerged as indispensable mediators of mechanosensing in both mammalian and plant cells. In mammalian cells, mechanoactivation was shown to trigger reorganization of the microtubule cytoskeleton, which is required to activate YAP/TAZ and Hippo signalling (75). Furthermore, rapid transduction — on the timescale of seconds and over distances of ∼10–20 μm — of localized oscillatory mechanical stress, as measured by Src kinase mechanical activation, was found to depend on and colocalize with sites of large microtubule displacement in 79% of cases (76). In plant cells, changes in anisotropic mechanical stress cause cortical microtubules to reorient along the direction of maximal tension, thereby guiding cellulose synthase complexes and reinforcing the cell wall (77). Notably, experiments from the same study suggest that microtubules in pavement cells experiencing tension may be resistant to katanin-mediated severing. Microtubules have therefore been proposed to act as tension sensors regulating anisotropic growth and tissue morphogenesis in plants; however, the molecular mechanisms underlying this mechanosensing remain largely unknown and have been identified as one of the major open questions in plant biology (78, 79).

Our data provide direct evidence that physiologically relevant tensile forces of less than 23 pN can induce microtubule lattice expansion and associated conformational changes. This suggests that mechanical force represents an additional layer of biochemical regulation of the microtubule interactome in cells, alongside tubulin isoform composition and post-translational modifications. This mechanochemical transduction, whereby changes in lattice conformation directly modulate biochemical interactions, occurs on a second timescale. We therefore propose that microtubules function as “communication lines”, transducing mechanical signals into biochemical regulation rapidly and with spatial precision throughout the cell body.

## Limitations

The assay developed in this work opens the way to address the limitations of this work by future studies to further investigate how microtubule mechanosensitivity is modulated by tubulin isoform diversity, different nucleotide analogues, and stabilizing reagents.

## Contributions

S.P. conceived the project. Y.L., B.F., J.M., E.S., E.M.O., N.M.R., I.K., and S.P. conceptualized research and designed the experiments. Y.L. and S.P. conducted the optical tweezers experiments. Y.L. and B.F. designed, expressed, and purified the kinesin-1 constructs. J.M. performed the cell experiments. S.P. and L.M. wrote software for data formatting and analysis. Y.L., B.F., S.P., and I.K. analysed fluorescence movies and kymographs. S.P. and I.K. analysed the optical tweezers data. I.K. developed the theoretical model and performed the simulations. All authors contributed to writing the manuscript and provided feedback.

## Acknowledgments

S.P. thanks Sven zur Oven Krockhaus (University of Tübingen) and Ole Schwarz (MPI Heidelberg) for insightful discussions on super-resolution localisation analysis of QDs. S.P. thanks Carolina Carrasco Pulido for useful discussions while establishing the single-bead assay with surface-tether microtubules. S.P. thanks Alan Boka, Daniel Safer, Mariko Tokito, Tanya Perez and Cece Petruconis for their technical support during the experiments at University of Pennsylvania. S.P. also thanks the laboratory of Dave Thirumalai (University of Texas at Austin) for stimulating discussions during a 10-day visit in 2019. The visit was supported by the Centre for Engineering MechanoBiology (CEMB), an NSF Science and Technology Centre, under grant agreement CMMI: 15-48571, while S.P. was working at the University of Pennsylvania. I.K. thanks the Leibniz Institute of Polymer Research in Dresden for hosting him as an invited scholar. N.M.R. and J.M. acknowledge support from the Deutsche Forschungsgemeinschaft (DFG, German Research Foundation; project number 335549539—GRK 2381) and core funding from the Excellence Strategy at the University of Tübingen. YL and BF acknowledge financial support by the DFG through the RTG 2364 “MOMbrane” and the University of Tübingen. S.P. acknowledges financial support by DFG “Eigene Stelle” grant (project number: PY 121/2-1).

## Materials and Methods

### Reagents

Streptavidin-conjugated quantum dots (QD585) were purchased from Thermo Fisher Scientific. Unlabelled, biotinylated, Rhodamine- and Alexa 488-labelled tubulins were prepared in-house from porcine brain (80) and also purchased from Cytoskeleton Inc. (Denver, CO, USA). Digoxigenin-labelled tubulin was prepared in-house (Supporting Information). NHS-Rhodamine (46406) and NHS-Alexa 488 (A20000) were purchased from ThermoFisher Scientific. NHS-Digoxigenin (11333054001) and Biotin-NHS (203112) were purchased from Merck. Sterile collodion (2% nitrocellulose in amyl acetate) and amyl acetate were purchased from Electron Microscopy Sciences (Hatfield, PA, USA) and from Merck (09817, W504009). Glass coverslips were sourced from multiple suppliers: 22 × 40 mm, No. 1.5 (ThermoFisher Scientific); 22 × 22 mm, 0.170 ± 0.005 mm, No. 1.5 (Marienfeld, Germany); and 18 × 18 mm, 0.170 ± 0.005 mm, No. 1.5 (Zeiss, Germany). Double-sided adhesive tape 3M 9832+ was purchased from ULINE, USA and from Klebershop24 GmbH & Co. (KS DC-3702-9, Koblenz, Germany). GMPCPP (guanosine-5′-[(α,β)-methylene]triphosphate, sodium salt) was purchased from Jena Bioscience (NU-405L, Jena, Germany). Paclitaxel was purchased from Merck, Germany (PHL89806) and Cytoskeleton Inc., USA (TXC01). ATP was purchased from Merck, Germany, (A2383). High-vacuum grease was purchased from Dow Corning (Midland, MI, USA). Trolox was purchased from Merck (648471). Neutravidin was purchased from ThermoFischer (31000). Casein from bovine milk was purchased from Merck. Sheep anti-digoxigenin antibody was purchased from Bio-Rad (3210-0488). D-Glucose, glucose oxidase from *Aspergillus niger* (G7141), and catalase from bovine liver (C1345) were obtained as lyophilised powders from Merck. Non-functionalised polystyrene microspheres (nominal diameter 2r = 0.8 µm, PS03004) were purchased from Bangs Laboratories (Fishers, IN, USA). Two-component epoxy minute adhesive was manufactured by Weicon and purchased by Conrad, Germany.

### Protein constructs

The protein constructs used in this study were the truncated (amino acids 1–430) kinesin-1 heavy chain isoform 5C from *Rattus norvegicus* rKIF5C(430) and the truncated (amino acids 1–560) kinesin-1 heavy chain isoform 5B from *Homo sapiens* hKIF5B(560). Fluorescent tags were fused to the C-terminus of both constructs. For more details about protein expression and purification see *Supporting Information*.

### Cell culture, plasmids and transfection

HeLaM cells were cultured in Dulbecco’s Modified Eagle Medium (DMEM; Thermo Fisher Scientific) supplemented with 10% fetal bovine serum (FBS; Thermo Fisher Scientific) and 1% penicillin–streptomycin (Thermo Fisher Scientific). Cells were maintained at 37 °C in a humidified incubator with 5% CO₂. eGFP–tau and mCherry–ensconsin constructs were obtained from Addgene (plasmids #46904 and #26742, respectively). For DNA transfection, cells were transfected using Lipofectamine™ 2000 (Invitrogen) according to the manufacturer’s instructions. Briefly, 1 µl of Lipofectamine 2000 was used per transfection with the indicated plasmids, and cells were imaged 24–48 h post-transfection.

### Cell Imaging and fluorescent microscopy

For all cellular imaging, cells were seeded onto glass-bottom MatTek dishes (MatTek Corporation) or ibidi dishes (ibidi GmbH). Imaging was performed using a Zeiss LSM 980 laser-scanning confocal microscope equipped with Airyscan 2 for enhanced resolution and sensitivity. Excitation was provided by diode and solid-state lasers spanning 405–640 nm, selected according to the fluorophores used. Emission signals were detected using GaAsP detectors in combination with Airyscan processing when applicable. Images were acquired using a Plan-Apochromat 60× oil-immersion objective (NA 1.45) and controlled using ZEN Blue software (Zeiss).

### Neutravidin coated bead preparation

Neutravidin-coated polystyrene beads were prepared by mixing 5 μL (1 μL) of 2% (10%) solid polystyrene beads 2r = 0.99 μm (2r = 0.80 μm) with Neutravidin to a final concentration of 6.5 g/L in a final volume of 31 μL (15 μL) in BRB80 (pH 6.9). The solution was incubated overnight at 4°C or for 2 hours at room temperature on a rotary wheel. Beads were then washed 10 times with 100 μL of BRB80 (pH 6.9) by centrifugation at 14,000 × g for 1.5 min at 4°C. After the final wash, beads were resuspended in 50 μL of 5 g/L casein and used within 12 hours.

### Preparation of stabilized microtubule and tubulin GMPCCP seeds

Doubly cycled GMPCPP-stabilized tubulin seeds were prepared as previously described (81). The final solution had a nominal tubulin concentration of 20 μM and was aliquoted into 3 μL portions, flash-frozen in liquid nitrogen, and stored at −80°C. Prior to use, aliquots were rapidly thawed by placing them on a pre-warmed aluminum block at 37°C, after which 8 μL of BRB80 pH 6.9 was added. Thawed aliquots were kept at room temperature, protected from light, and used within 3 days.

Doubly stabilized microtubules for the dual laser-trapped experiments were prepared using GMPCPP and paclitaxel as follows. A total tubulin concentration of 40 μM (43% biotinylated tubulin, 12 % rhodamine-tubulin) was polymerized in the presence of 1 mM GMPCPP in BRB80 (pH 6.9) at 37°C for 1–2 hours. Polymerized microtubules were collected by centrifugation at 20,000 × g for 25 min at 25°C. The supernatant was discarded and the pellet was gently washed twice with 40 μL of warm BRB80 before being resuspended in 20 μL of BRB80 (pH 6.9) supplemented with 40 μM paclitaxel.

### Optical Tweezers Experiments

For preparation of the experimental chambers see *Supporting Information*. The dual-trap experiments were performed at E. Mike Ostap’s lab at the University of Pennsylvania (for setup see *Supporting Information*). For the dual-trap experiments, GMPCPP- and paclitaxel-stabilized microtubules (50% biotinylated tubulin) were used to form dumbbells as previously described (29), using In house neutravidin-coated PS beads 2r = 1.00 um (07310, Polysciences, PA, USA). Microtubules were decorated with streptavidin-conjugated QDs 585 by mixing 0.66–5 nM QDs with 250 nM microtubules supplemented with 20 uM paclitaxel, allowing the mixture to incubate for at least 2 min prior to diluting 1 to 5 in final solution containing 20 μM paclitaxel, 2 g/L casein, 42 mM D-glucose, 190 U/mL glucose oxidase, and 0.2 U/mL catalase, and injection into the flow cell. The open ends of the flow cell were then sealed with vacuum grease. BRB80 (pH 7.5) was used in these experiments, as the optimal pH range for photostability of the QDs is between 7 and 8. Both traps were of approximately equal stiffness. Trap stiffness and the voltage-to-force conversion factor were determined by fitting the power spectrum of the Brownian motion of a trapped bead to a Lorentzian function. Tensile force on the dumbbell was controlled by displacing one of the two traps by a defined distance using an acousto-optic deflector.

The single-trap experiments were performed at Erik Schaffer’s lab at the University of Tubingen (for setup see *Supporting Information*). For the single-trap experiments neutravidin-coated PS beads 2r = 0.784 um were used. Flow cells were first washed with 100 μL of 5 g/L casein supplemented with 20 μM paclitaxel, prior to injection of the experimental solution consisting of: 20 mM D-glucose, 0.008mg/mL glucose oxidase, 0.02 mg/mL catalase, 20 μM paclitaxel, 3 mM Trolox, 1 mM DTT, 5 g/L casein, 2 mM MgATP, 0.64–1.1 nM kinesin motor, and Neutravidin-coated polystyrene beads diluted 1:20–1:60 from the washed bead stock (see “Neutravidin coated bead preparation”). Trap stiffness and the voltage-to-force conversion factor were determined as described above for the x, y and z directions. To maintain a constant tensile force, a closed-loop feedback system was implemented in which the QPD signal served as the error signal. The resulting correction voltage was amplified and applied to a piezoelectric stage, which adjusted its position to minimize the error and maintain the desired force level. The force setpoint was updated manually at approximately one-minute intervals.

### Fluorescence Imaging

For the dual-trap experiments, fluorescence imaging of QDs was performed using an EMCCD camera (iXon3 897, DU-897E-CS0-#BV, Andor Technology, Belfast, UK) with the following acquisition parameters: sensor temperature −80°C, exposure time 100 ms, frame rate 9.04 Hz, EM gain 200, 1 × 1 binning (pixel size 46 nm). Illumination was provided by a 488 nm laser (Coherent, Santa Clara, CA, USA) at 10% power in epifluorescence mode.

For single-trap experiments, fluorescence imaging was performed using an sCMOS camera (ORCA-Flash4.0, C11440, Hamamatsu, Japan) with the following acquisition parameters: sensor temperature −20°C, exposure time 200 ms (continuous acquisition), 2×2 binning (pixel size 148 nm). Illumination was provided by a 488 nm, 80 mW diode laser (PhoxX® 488-80, Omicron-Laserage Laserprodukte GmbH, Rodgau, Germany) and a 561 nm, 50 mW laser (OBIS 561 LS, Coherent, Santa Clara, CA, USA), each operated at 5% power in HILO illumination mode. Kinesin molecules were imaged under 488 nm excitation and surface-tethered microtubules under 561 nm excitation. To minimize photobleaching and photodamage, both laser beams were further attenuated using a 5% neutral density filter placed in the illumination path upstream of the objective.

### Lattice and Motor Simulations

The tubulin heterodimer at each section is assumed to be switchable between two discrete states ±1 and is harboring kinesin motors that are binding/unbinding and running while responding to the lattice state. Simulated kinesins bind only to the short state,−1, while they are forbidden from binding to the long-state, +1. Each section of the microtubule lattice follows an Ising-Glauber type dynamics (82) of the two states ±1, with a mechano-chemical biasing field *H*, and a coupling energy between subsequent sections *J*. A homogeneous microtubule lattice is characterized by the two Ising energetic parameters B and J and a single dynamic parameter, the attempt frequency of lattice section flipping f, determining with J and H the rate of domain creation and growth respectively. A non-homogeneous, patchy microtubule lattice is modeled by introducing a position dependent local field H_b_(i) biasing at the site i the section state energetically towards one of the two states. The kinesins are characterized by their association and dissociation kinetic rates and their running speed and are randomly, asynchronously positioned and state actuated over the simulation cycle followed by an asynchronous Glauber actuation of the tubulin lattice states. The microtubule simulation parameters are chosen in agreement with the critical force extracted from pulling data (Figure 1 E). The kinesin motor number, binding, unbinding rates and running speeds are chosen in agreement with experimental motor speeds, densities and lifetimes (cf. Fig. 4 caption).

### Data Analysis

Data analysis was done using software Origin2026 and Python. For more details on the specific analysis of the various experimental datasets see *Supporting Information*.

### Language Editing

Claude Sonnet 4.6 (Anthropic) was used for assistance with manuscript editing and writing.

