## Supplemental Information for "Mechanical tension expands the microtubule lattice stepwise and modulates kinesin-1 binding in an isoform-dependent manner"

### Supporting Materials and Methods

#### Protein expression of rKIF5C(430)

The pET17-b (MilliporeSigma, Novagen) plasmid of the truncated (amino acids 1–430) kinesin-1 heavy chain isoform 5C from *Rattus norvegicus* rKIF5C(430) used in this work is based on a plasmid originally provided by the Howard Laboratory (Yale University, New Haven, CT). The rKIF5C(430) construct is expressed with a C-terminal GFP and a hexahistidine-tag. First, the plasmid was transformed into Rosetta 2 (DE3, *E. coli*) via heat shock and plated on LB agar plates (lysogeny broth) containing 100 µg/mL ampicillin and 25 µg/mL chloramphenicol. A single colony was used to prepare a 12 h starting culture. Cells were further grown in phosphate-buffered TB medium (terrific broth) supplemented with 89 mM sodium phosphate buffer (pH 7.5), 1 mM glucose, 1 mM $MgCl_2$, 100 µg/ml ampicillin, 25 µg/ml chloramphenicol at 37°C and 250 rpm until an $OD_{600}$ of 0.6 and cooled to 18°C before protein expression was induced with 1 mM IPTG (isopropyl β-D-1-thiogalactopyranoside) for 16 h. Bacterial cells were harvested by centrifugation, the cell pellet was flash frozen via liquid nitrogen and stored at -80°C.

#### Protein expression of hKIF5B(560)

The plasmid encoding the truncated (amino acids 1–560) kinesin-1 heavy chain isoform 5B from *Homo sapiens* (UniProtKB accession P33176) with a C-terminal mNeonGreen and an octahistidine-tag used in this work was custom synthesized by GenScript (Piscataway, NJ, USA). The plasmid was expressed with the baculovirus based expression ExpiSf Expression System (Gibco, ThermoFisher Scientific), which uses ExpiSf cells (Gibco,A35243, derivative of Sf9 *Spodoptera frugiperda*) To generate bacmid, the plasmid was transformed into DH10Bac cells (ThermoFisher,10361012) via heat shock and plated on LB agar plates (lysogeny broth) containing 50 µg/mL kanamycin, 7 µg/mL gentamycin, 10 µg/mL tetracyclin, 20 µg/mL X-Gal and 40 µg/mL IPTG. Positive clones were distinguished via two rounds of blue-white screening and colony PCR. A single colony was used to inoculate 30mL of a LB-media containing 50 µg/mL kanamycin, 7 µg/mL gentamycin,10 µg/mL tetracyclin and incubated for 12 h o/n at 250 rpm and 37 °C. The next day bacmid was purified via the Pure Link HiPure Plasmid DNA Purification Kit (Invitrogen, K2100-01) following the standard protocol. To generate $P_0$ Baculovirus, a running culture of ExpiSF cells where transfected with the prepared baculovirus following the ExpiSF standard protocol for $P_0$-preparation in 25 mL cultures. The $P_0$ virus had a sufficient enough titer to successfully infect insect cell cultures and was thus used directly for the infection and protein expression. The protein was expressed in fresh ExpiSf cultures following the standard ExpiSF expression protocol. Insect cells were harvested after three days by centrifugation. The pellet was washed by resuspension in PBS buffer. The cell pellet was then flash frozen via liquid nitrogen and stored at -80°C.

**Protein purification of rKIF5C(430) and hKIF5B(560)**

Protein purification was carried out at 4°C or on ice. Buffers were adjusted to a final pH of 7.5 The cell pellet was thawed and resuspended in lysis buffer containing 50 mM sodium phosphate (pH 7.5), 300 mM KCl, 1 mM $MgCl_2$, 20 mM imidazole, 5% glycerol, 0.1% Tween-20, 0.1 mM ATP, 5 mM β-mercaptoethanol (β-ME), 1 mM PMSF, 25 U/mL benzonase, and 1× cOmplete EDTA-free protease inhibitor cocktail (Roche). The resuspended pellet was lysed using a tip sonicator (Active Motif, Q120AM) at 10 W for 3 min with a pulse cycle of 3 s on and 7 s off. The lysate was clarified by two rounds of centrifugation at 7200 g for 20 min and 30000 g for 30 min. The resulting supernatant was used for immobilized metal affinity chromatography (IMAC) and loaded onto a 5 mL HiTrap TALON crude column (Cytiva, 28953767). After washing with washing buffer (50 mM sodium phosphate (pH 7.5),, 300 mM KCl, 1 mM $MgCl_2$, 60 mM imidazole, 10% glycerol, 0.1 mM ATP, 5 mM β-ME, and 1× cOmplete EDTA-free protease inhibitor cocktail), the protein was eluted with elution buffer containing 50 mM sodium phosphate (pH 7.5), 300 mM KCl, 1 mM $MgCl_2$, 500 mM imidazole, 10% glycerol, 0.1 mM ATP, 5 mM β-ME, and 1× cOmplete EDTA-free protease inhibitor cocktail. The eluted protein was then loaded onto a HiTrap Desalting column (Cytiva, 29048684) and eluted with cleanup buffer consisting of 50 mM sodium phosphate (pH 7.5), 300 mM KCl, 1 mM $MgCl_2$, 10% glycerol, 0.1 mM ATP, 5 mM β-ME, and 1× cOmplete EDTA-free protease inhibitor cocktail. Finally, the protein was, aliquoted, snap-frozen in liquid nitrogen, and stored at −80 °C.

**Preparation of Experimental Flow Cells**

For the dual-trap laser tweezer experiments using microtubule dumbbells decorated with QDs, chambers were prepared as previously described (1), with the only difference that double-sided sticky tape 3M 9832+, ( ULINE, USA) was used instead of that previously reported. The bottom coverslip of the chamber was sparsely decorated with immobilized spherical silica pedestals (2r = 2.5 µm; SS05000, Bangs Labs, USA) to allow identification of the bottom surface under transmission light illumination.

For the single-trap experiments, 22 × 22 mm and 18 × 18 mm glass coverslips coated with 0.1% collodion (nitrocellulose) were used to assemble the flow chamber as follows. The 22 × 22 mm coverslip was placed on the bottom, and double-sided sticky tape was applied along two parallel sides to define the flow channel. Two gel-loading tips were placed narrow-end down, one on each side of the flow channel and parallel to the tape, serving as inflow and outflow tubing after sealing. The 18 × 18 mm coverslip was then placed on top, and two-component one-minute epoxy was applied around the entire perimeter of the top coverslip, including the flow channel openings, to prevent leakage. Assembled chambers were left overnight to allow complete curing of the epoxy and were used the following day. The flow channel volume was ≤ 10 µL. All solutions were prepared in BRB80 (pH 6.9) and introduced into the chamber by pipetting directly through the wide opening of the gel-loading tips in the following sequence: 25 µL of 0.1 g/L anti-digoxigenin antibody (5 min incubation), 50 µL of 5 g/L casein (10 min incubation), 60 µL of 17–68 nM GMPCPP-stabilized seeds (20% digoxigenin-tubulin, 14% rhodamine-tubulin; flow-through only, no incubation), followed by 80 µL of 5 g/L casein as a wash step.

Subsequently, 25 µL of elongation mix was injected, consisting of 14 µM tubulin (2% rhodamine-tubulin), 1 mM GMPCPP, and 5 g/L casein. The inflow and outflow openings were sealed with Parafilm to prevent evaporation, and the flow cell was placed in a humidity chamber on a hot plate (Premiere XH-2002 Slides Warmer, Microscopes America, Inc, Georgia, US) at 37°C for 1.5–2 hours, protected from light. The flow cell was then washed with 100 µL of 5 g/L casein, after which 25 µL of "capping" solution was injected, consisting of 14 µM tubulin (10% biotinylated tubulin, 14% rhodamine-tubulin), 1 mM GMPCPP, and 5 g/L casein. The inflow and outflow openings were again sealed with Parafilm, and the flow cell was incubated in a humidity chamber on a hot plate at 37°C for 1 hour. Finally, the flow cell was washed with 100 µL of 5 g/L casein and 4 g/L BSA supplemented with 40 µM paclitaxel. Flow cells were stored at room temperature in a humidity chamber protected from light and used no earlier than 2 hours and no later than 18 hours after preparation.

**Optical Tweezers Setups**

The dual-trap custom-built optical tweezer setup at the Ostap lab (University of Pennsylvania) has been previously described in detail (2). Briefly, the system consists of a dual-beam optical trap using a 63× water-immersion objective (C-Apochromat, Zeiss, Jena, Germany) with a numerical aperture of 1.2. The trap stiffness (pN/nm) and the voltage-to-force conversion factor (pN/V) were determined for each trapped bead prior to microtubule attachment by fitting a Lorentzian function to the power spectrum of the Brownian motion of the trapped bead. Tensile force was applied to microtubule dumbbells by increasing the distance between the two laser foci, achieved by displacing one of the two traps using an acousto-optic deflector. The stiffness of the individual laser traps was in the range of 0.04 – 0.11 pN/nm. Data were recorded at 19°C, low-pass filtered at 1 kHz, digitized at a sampling rate of 2 kHz, and acquired using custom software written in LabVIEW (National Instruments, Austin, TX, USA).

For all QD experiments, a 488 nm laser was used to illuminate the sample in epi-fluorescence mode from the condenser side, with the excitation beam entering through the top coverslip of the experimental chamber. The instrument is based on an upright modified Olympus microscope, in which the trapping laser is focused through the imaging objective from below, entering through the bottom coverslip of the experimental chamber. The same objective was used for both optical trapping and fluorescence image collection.

The custom-built single-trap optical tweezers setup at the Schäffer lab (University of Tübingen) uses a 1064 nm continuous-wave laser (3 W, diode-pumped Nd:$YVO_4$, Smart Laser Systems, Germany) for optical trapping, steerable in the image plane via a piezo tip-tilt mirror (PTM, TT2.5, Piezoconcept, France). For detection, the back focal plane of the condenser lens (D-CUO Achr.-Apl. NA 1.4, Nikon Instruments, Japan) was imaged onto a quadrant photo diode (QP154-Q-HVSD, First Sensor AG, Germany) for back focal plane interferometry. For sample positioning, a piezo translation stage was used (NPXY100Z50-244 with a digital controller LC.403 3-CH, DSP Npoint, USA) having a travel range of 100 µm and 50 µm for the lateral and vertical axes, respectively. For coarse positioning, another stage was mounted on top of the latter piezo translation stage (12 mm travel range, 3x SLC1720, SmarAct, Germany). The setup is controlled by custom written LabVIEW (National Instruments, Austin, USA) programs. Data were recorded at 25°C and 100 kHz.

The setup is coupled to total internal reflection fluorescence (TIRF) and interference reflection microscopy (IRM) (3, 4). Fluorescence excitation lasers (488 nm and 561 nm) were introduced via micromirrors positioned directly beneath the objective (CFI Apo TIRF 60×, oil immersion, NA 1.49, Nikon Instruments), and the emission was spectrally split and projected onto an sCMOS camera.

**Control Experiment with Surface immobilized QDs**:
A nitrocellulose-coated flow cell was incubated with a $1{:}25 \times 10^4$ dilution of QD stock solution in BRB80 (pH 7.5) for 1 min, then washed extensively (~10 chamber volumes) with BRB80 (pH 7.5) supplemented with glucose and an oxygen-scavenging system (glucose oxidase and catalase, GOC). Both ends of the chamber were sealed to prevent evaporation. To verify that the microscopy setup and imaging conditions provided sufficient precision for super-resolution localization of QDs, we first performed control experiments with surface-immobilized QDs. Their mean positions over 300 frames could be determined with a standard error of the mean below 3 nm (Fig. S1). Upon stage displacements the change in the mean positions of the QDs across the field of view were tightly clustered around the same value with standard deviation less than 3 nm as well (Fig. S1).

**Data Analysis.**

**Assumption of strictly elastic deformation of microtubule lattice for F< 23 pN**

The absence of a linear elastic response, although surprising at first glance, can be justified on simple geometric and physical grounds. Assuming hypothetically that the measured expansion of the microtubules would be strictly elastic, the Young's modulus $E$ can be estimated as $E = \langle \Delta F \rangle / (\varepsilon \times A) = \Delta\sigma/\varepsilon$, where $\varepsilon = 0.0033$, $\langle \Delta F \rangle = 10$ pN, $\Delta\sigma = \langle \Delta F \rangle / A$ is the change in stress associated with the force change $\langle \Delta F \rangle$, and $A = 314$ nm² is the cross-sectional area of a hollow cylinder with inner and outer radii of 7.5 nm and 12.5 nm, respectively (5, 6). This estimate would yield a Young's modulus of ~11 MPa, which is more than two orders of magnitude smaller than the expected elastic moduli of ~ 1 GPa for the tubulin lattice ((7) and citations therein). An alternative explanation for this anomalously soft modulus is that the microtubule mechanical response originates from inter-protofilament shear displacement. This interpretation, however, appears highly improbable on physical grounds: for a microtubule of typical length ~15 µm, a 0.33% elongation corresponds to a net displacement of ~50 nm — equivalent to ~6 tubulin heterodimer lengths — implying a dramatic register shift between neighbouring protofilaments. Such a large-scale lattice shear would transiently rupture lateral contacts across the entire length of the microtubule and would most likely result in lattice disintegration. Consistently, cryo-electron tomograms of both straight and highly bent or flattened microtubules reveal no significant inter-protofilament shear displacement, with lateral shifts confined to the range −0.5 to +0.9 Å (8). Structural and fluorescence data combined (9–12) suggest that the dominant contribution to microtubule lattice compaction and expansion is not inter-protofilament shear but rather a change in the axial length of the tubulin heterodimer itself.

**Super-resolution tracking**

For each dumbbell, the QD with the highest positional precision was chosen as the reference point, relative to which the distances $L$ of the other QDs along the microtubule lattice were measured. Super-resolution localization of QDs was performed using the software ThunderSTORM (13). The theoretically expected standard deviation ($\sigma$) of the point spread function (PSF) for an emission wavelength of $\lambda$ = 585 nm and a numerical aperture of NA = 1.2 is given by the approximation for a 2D Gaussian PSF: $\sigma \approx 0.21 \cdot \lambda/\mathrm{NA}$ = 102.4 nm (Ref). Localizations were filtered on two criteria: the fitted PSF width $\sigma$ had to exceed this theoretical lower bound, and the localization uncertainty $\Delta x$ had to be less than 20 nm. Additionally, only QDs whose x and y localization distributions were each well described by a symmetric Gaussian were retained for further analysis. For each qualifying QD, the mean x and y coordinates were determined, and the distance L between pairs of QDs was calculated from these means. The associated uncertainty $\sigma_L$ was then obtained by standard propagation of the corresponding standard errors. For each microtubule dumbbell, all pairwise distances between QDs were calculated relative to the QD with the smallest localization error, thereby minimizing the cumulative error in the distance calculation.
To validate the reliability of our super-resolution localization approach, we performed a control experiment in which surface-immobilized QDs were imaged at two piezo stage positions separated by a known displacement of 100 nm along the x-axis (Fig. S1). The x and y position coordinates of 15 QDs tracked across the field of view followed 2D Gaussian distributions, as expected for well-localized single emitters (Fig.

S1D). The average measured displacement across all 15 QDs was 88.7 nm with a standard deviation of 2.24 nm (2.5%). The small but systematic deviation from the nominal 100 nm displacement, as well as the presence of a minor off-axis component, are attributable to the fact that the coverslip is not rigidly clamped to the stage, permitting slight relative sliding between the flow cell and the piezo stage.

For super-resolution localization of QDs attached to microtubule dumbbells, only QDs located more than 1 μm from the polystyrene beads were included in the analysis. This is because the microtubule region in close proximity to the beads may undergo changes in curvature upon changes in applied force, as the beads rotate in response to changes in the associated torques.

The finite positional resolution $\sigma_{\Delta L}$ of the relative distances $\Delta L$ between pairs of QDs imposes an L-dependent limit on the accurate determination of strain ε, since $\Delta L = \Delta\varepsilon \cdot L_{\mathrm{init}} \geq \sigma_{\Delta L}$, where $L_{\mathrm{init}}$ is the initial distance between the QD pair. Specifically, for a given magnitude of strain ε, reliable strain measurements require $L_{\mathrm{init}} \geq \sigma_{\Delta L}/\Delta\varepsilon = L_{\mathrm{threshold}}$. The slope of $\Delta L$ as a function of $L_{\mathrm{init}}$ for all Qd pairs upon decrease ($\Delta F < 0$) and increase ($\Delta F > 0$) of the tensile force F applied to the microtubule dumbbells yields the corresponding strain values $\Delta\varepsilon_{\Delta F<0} = (0.17 \pm 0.16)\%$ and $\Delta\varepsilon_{\Delta F>0} = (0.26 \pm 0.14)\%$ (95% CI) (Fig. S2A). Assuming an average strain of $\langle\varepsilon\rangle = 0.0022$, and given that the standard error of the mean QD positions does not exceed 3 nm, application of the error propagation formula successively for $\sigma_L$ and then $\sigma_{\Delta L}$ yields $\sigma_{\Delta L} = 2 \times 3$ nm = 6 nm. The expected $L_{\mathrm{threshold}}$ is therefore 6 nm/0.0022 ≈ 2727 nm, consistent with the observed behaviour in the plot of change in strain $\Delta\varepsilon$ versus $L_{\mathrm{init}}$ for all QD pairs, where the scatter in $\Delta\varepsilon$ values is markedly inflated below $L_{\mathrm{init}} \sim 2000$ nm, for both $\Delta F < 0$ and $\Delta F > 0$ (Fig. S2B).

To determine $L_{\mathrm{threshold}}$ more accurately directly from the data, the average strain $\langle\Delta\varepsilon(L_{\mathrm{init}} > \min(L_{\mathrm{init}}))\rangle$ was calculated by serially and cumulatively excluding the smallest $L_{\mathrm{init}}$ value at each iteration, until the remaining number of data points was no fewer than 30, to ensure robust statistics (Fig. S2C). The standard error of the mean strain ε is expected to initially decrease as the noisiest short-distance pairs are excluded, but to subsequently increase as the sample size is reduced, for both $\Delta F < 0$ and $\Delta F > 0$ (Fig. S2D). Of the two values of $L_{\mathrm{init}}$ that minimize the SEM of ε for $\Delta F < 0$ and $\Delta F > 0$ respectively, the larger value (2500 nm) was selected as $L_{\mathrm{threshold}}$ to filter the data presented in Fig. 1D of the main text.

It is worth mentioning that similar behaviour of strain ε versus $L_{init}$ is observed for surface immobilized QDs (Fig. S5)

**Analysis of fluorescent movies, kymograph quantification and running motor density profile**

Surface-immobilised microtubules or bright fluorescent spots were used as fiducial markers to identify stage motion corresponding to changes in the tensile force applied to the microtubule to which the laser-trapped, neutravidin-coated bead was stably attached. Based on this, each movie was segmented into sub-movies corresponding to distinct constant force regimes. Maximum intensity projections were calculated for both excitation channels (561 nm and 488 nm). Tracks were identified manually using ImageJ. Only tracks longer than 3 pixels were retained. In the rare cases where two tracks merged, both were discarded from the analysis. Run lengths were calculated only within the segment of the microtubule between the seed and the cap. The cap and seed regions were excluded due to the high percentage of fluorescently labelled tubulin (12%), which was found to produce occasional artefacts such as the termination of run lengths at a fixed position.

The running motor density profiles were calculated via bootstrapping of the track set from the corresponding kymographs. For each bootstrap replicate, the average projection of the tracks along the microtubule axis was calculated using a fixed bin size of 4 pixels (592 nm). The 95% CI of the mean across bootstrap replicates represents the statistical uncertainty of the density profile. During bootstrapping, tracks were not shifted in time or space.

The variance of the running motor density profile was chosen as the statistical metric to assess whether all microtubule segments are equally accessible to motors. Under the null hypothesis of uniform motor accessibility, the density profile is expected to be uniform, which minimizes the variance for a given total number of motor events. Deviations from uniformity — arising from segments that preferentially attract or repel motors — produce a higher variance, making it a sensitive and natural measure of spatial heterogeneity. Pairwise comparisons between individual segments were avoided for two reasons: the small number of motors per segment reduces statistical power, and the large number of comparisons required substantially increases the probability of Type I errors (false positives).

To compare the variance of a given experimental running motor density profile with that expected for a uniform distribution along the microtubule, tracks were randomly offset along the microtubule axis during bootstrapping. When a track extended beyond the plus-end proximal boundary of the microtubule, the remainder was wrapped cyclically to the beginning of the microtubule. Not any spatial exclusion principle between the track was applied since more that one motor can exist in one pixel (pixel size 148 nm). This procedure produced a uniform mean distribution by construction. The 95% CI of the variance was then calculated from the distribution of bootstrap replicate variances, and the variance of the experimental profile was deemed statistically significant or non-significant relative to the uniform distribution accordingly. The p-value was calculated as the fraction of replicates with variance larger than the experimentally observed variance, i.e., $p = \frac{1+N(var_{bootstrap} \geq var_{obs})}{1+N_{tot}}$, where $N(var_{bootstrap} \geq var_{obs})$ represents the number of bootstrap replicates with variance larger or equal to the experimentally observed variance and $N_{tot}$ represents the total number of replicates. Bootstrapping and 95% confidence intervals were calculated using custom LabVIEW code. The two-tailed Mann-Whitney U test and data visualization were performed using OriginPro 2026. A two-tailed rather than one-tailed test was used, as the direction of the effect of force application on either QD interdistance or kinesin association and dissociation rates was not known a priori. All figures were prepared using the open-source software Inkscape.

**Relative chord elongation of a circular arc due to uniform change in curvature at constant arc length.**

To exclude the possibility that the increased QD separation, and therefore the measured change in strain $\Delta\varepsilon$, arises from force-induced global straightening of the microtubule (i.e., a uniform reduction in its overall, arc-like curvature) rather than from lattice expansion, we plotted the measured change in strain as a function of the initial distance between the pairs of QDs. If microtubule straightening were the dominant contribution, the apparent change in strain would be expected to increase with the initial separation between QDs (see calculation below and Fig. S4). However, we observed no statistically significant correlation between change in strain and initial QD spacing (Fig. S5).Let's assume an arc of initial curvature $R_i$ and initial angle $\theta_i < \pi$ (Fig. S4). Then the length of the cord connecting its two ends will be $L_i = R_i \cdot \sin\theta_i$. Upon application of extensile force, the curvature and the angle are changing to $R_f$ and $\theta_f$ (Fig. S4). The new cord length is $L_f = R_f \cdot \sin\theta_f$ but the length of the arc itself is not changing, i.e., $S = 2R_i \cdot \theta_i = 2R_f \cdot \theta_f$.

Then, the apparent strain $\varepsilon$ can be written as:

$\varepsilon = (L_f - L_i)/ L_i = L_f/L_i - 1 = R_f \cdot \sin\theta_f/R_i \cdot \sin\theta_i - 1 = \theta_i \cdot \sin\theta_f/\theta_f \cdot \sin\theta_i - 1 = (\sin\theta_f/\theta_f)/(\sin\theta_i/\theta_i) - 1$

For small angles approximation we apply Taylor's expansion around 0:

$\sin\theta \approx \theta - \theta^3/6$ and therefore $\sin\theta/\theta \approx 1 - \theta^2/6$

For small deformations $\theta_f$ can be expressed as $\theta_f = \theta_i - \delta$, where $\delta << 1$.

Applying the above approximations one gets:

$\varepsilon = [1 - (\theta_i - \delta)^2/6]/[1 - \theta_i^2/6] - 1 = [1 - (\theta_i^2 + \delta^2 - 2\theta_i\delta)/6]/[1 - \theta_i^2/6] - 1$

After applying the approximation $1/(1-x) \approx 1+x$ for $x<<1$ and keeping only first order terms of $\delta$ and $\theta_i$ one gets

$\varepsilon \approx \theta_i \cdot \delta/3$.

The strain therefore is proportional to the initial half angle and therefore the initial cord length $L_i$.

### Mapping of the Ising model from a magnetic to a polymorphic system

The polymorphic lattice model presented in the main text has a close analogy with the classical ferromagnetic 1D Ising model. The latter correspondence allows for the calculation of all relevant thermodynamic and statistical quantities, including the response of the strain to the external tension, Eq. (3) main text and the correlation length $\xi$. A useful summary of the 1D ferromagnetic Ising model is given in (14). The mapping between the polymorphic and ferromagnetic system is given in the following dictionary table ST1

**Table S1: Mapping magnetic to polymorphic system**

| Magnetic System | Polymorphic System | Replacement Rule |
|---|---|---|
| $S_i \in \{-1,1\}$ (spin) | $\sigma_i \in \{0,1\}$ (switch) | $\sigma_i \to \frac{S_i + 1}{2}$ |

| | | |
|---|---|---|
| $\mu$ (magn. moment per spin) | $\varepsilon_{sw}$ (transition strain) | $\varepsilon_{sw} \to \mu$ |
| $m = \frac{\mu}{N}\sum_{i=1}^{N}\langle S_i\rangle$ (mean magnet.) | $\varepsilon = \frac{\varepsilon_{sw}}{N}\sum_{i=1}^{N}\langle \sigma_i\rangle$ (mean strain) | $\varepsilon \to \frac{m+1}{2}$ |
| $m \in [-\mu, \mu]$ | $\varepsilon \in [0, \varepsilon_{sw}]$ | |
| $J$ (>0, ferromag. coupling) | $J$ (>0, allosteric coupling) | $J \to J$ |
| $H$ (ext. magnetic field) | $F + \frac{\Delta G}{l_{\text{lattice}}}$ (effective force) | $H \to F + \frac{\Delta G}{l_{\text{lattice}}}$ |
| $\xi$ (magn. correlation length) | $\xi$ (strain correlation length) | $\frac{\xi}{l_{\text{lattice}}} \sim \exp\frac{2J}{k_B T}$ (for $H \ll J$) |

Here the unit length of the lattice $l_{\text{lattice}} \approx 8nm$ for the MT case, $\Delta G$ the force-free free-energy bias of the short state vs. long. The correlation length $\xi$ has a more complex behavior for larger $H$, however simplifies to the classical expression for small ratio $H/J$ (~$10^{-2}$ here) given in the table.

### Supporting Figures

**Fig. S1:**

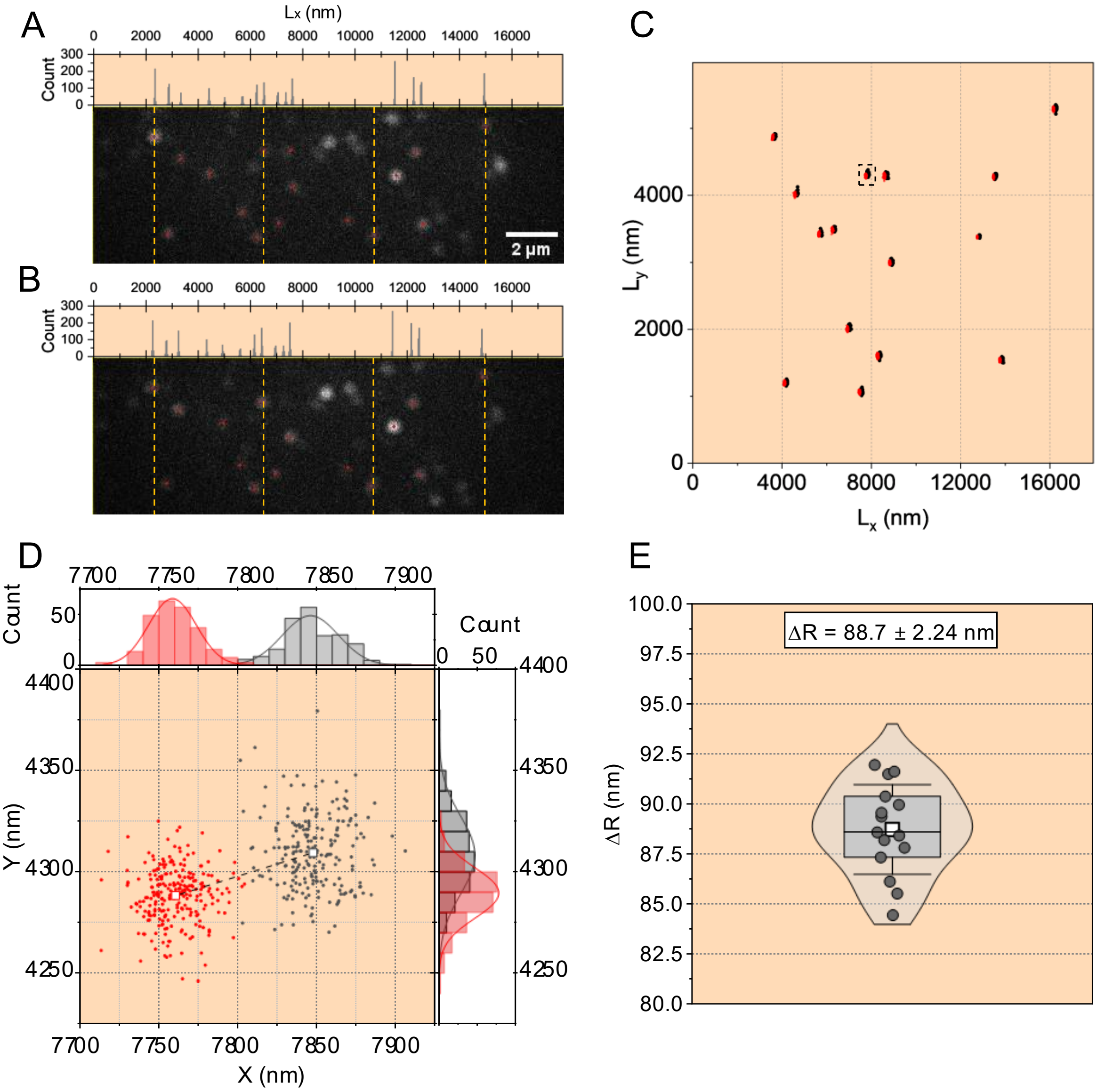


**Validation of super-resolution tracking of surface immobilized QDs.** (A) Single frame from a 300-frame movie showing surface-immobilized quantum dots (QDs) emitting at 585 nm. The centers of 15 randomly selected QDs are marked with red dots, and the vertical dashed yellow lines serve as guides to the eye. The histogram above the image shows the x-positions of these 15 QDs over all 300 frames, obtained using the ThunderSTORM localization software [9]. (B) Same as in (A), but with the piezo stage shifted by ~100 nm to the left along the x-axis. (C) Combined (x, y) localization positions of the QDs from the movies in (A) and (B), shown in black (●) and red (●), respectively. Because of the large plot scale, the dense localization clusters for individual QDs appear as single spots. The QD highlighted by the dashed black rectangle is analyzed further in (D). (D) Localization distribution of the QD marked in (C) for the movies corresponding to (A) (black; ●) and (B) (red; ●). Histograms of the x- and y-positions are shown above and to the right of the main plot, and solid curves represent Gaussian fits. The mean (x, y) positions for each condition are indicated by open squares of the corresponding color. The displacement ΔR between the two mean positions, shown by the dashed black line, reflects the stage shift. (E) Statistical violin-boxplot of the displacement ΔR for each QD, calculated from panel (C). The mean displacement across all traced QDs is $\langle \Delta R \rangle = 88.7 \pm 2.24$ nm. The narrow distribution of ΔR values demonstrates that the magnification and illumination are sufficiently uniform across the field of view and that ThunderSTORM provides reliable localization performance.

**Fig. S2:**

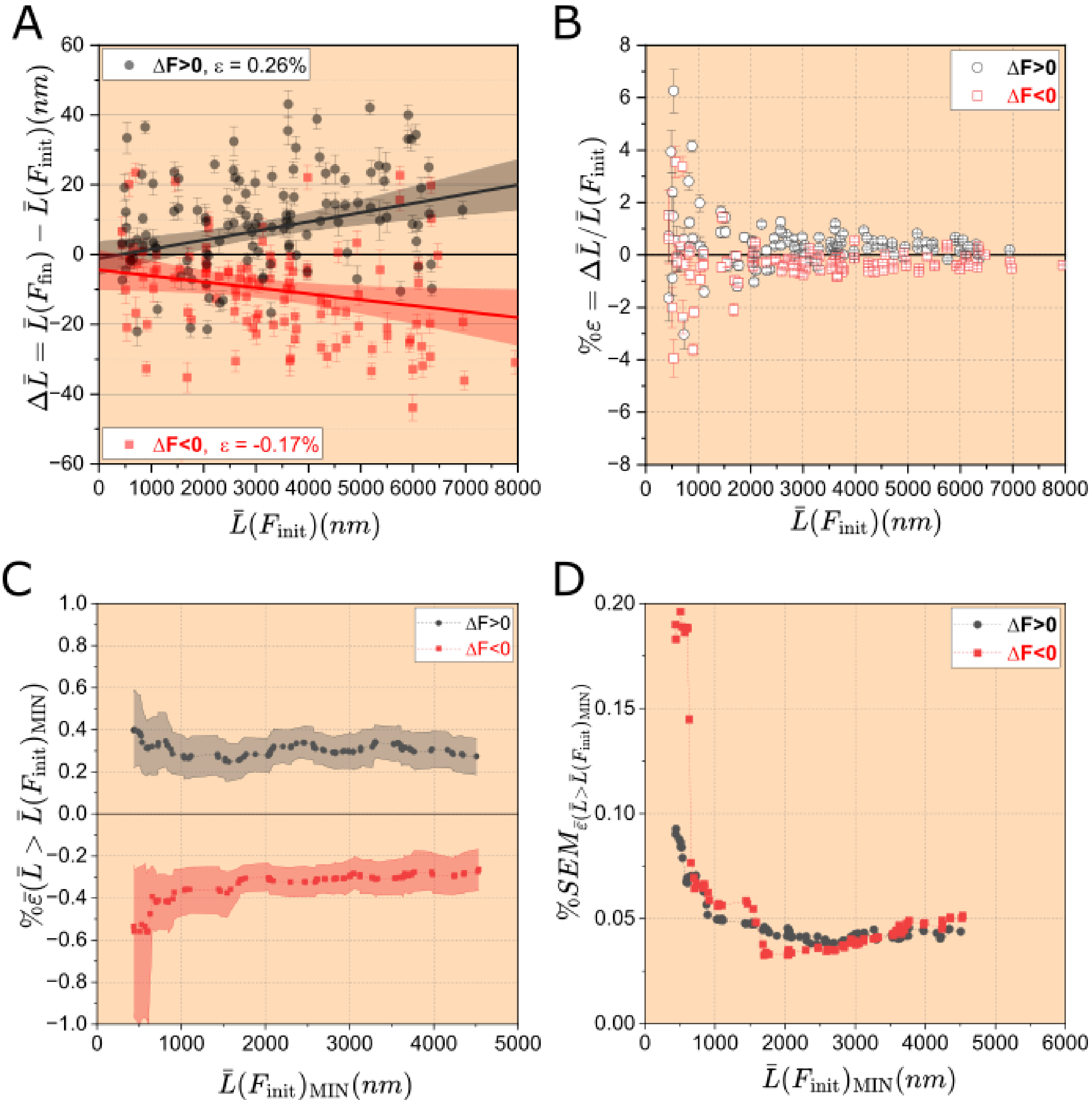


**L-dependent limit on strain calculation.** (A) Scatter plot showing the change in average distance ($\Delta\acute{L}$) between all pairs of quantum dots as a function of their average initial separation, $\acute{L}(F_{\text{init}})$, for $\Delta F > 0$ (black filled circles, N = 121) and $\Delta F < 0$ (red filled squares, N = 117). Error bars were calculated using standard error–propagation formulas based on the super-resolution positional tracking uncertainty of each quantum dot. Solid lines indicate linear fits to the data (color-coded accordingly), with shaded regions representing the 95% confidence intervals (CI) of the fits. The slope of each fit corresponds to the strain (ε) for the respective dataset, and the y-intercept is not significantly different from zero within the 95% CI. The absolute values of the fitted slopes overlap within their 95% CIs, indicating mechanical reversibility. (B) Strain

(%ε) plotted as a function of the average initial separation, $\acute{L}(F_{\text{init}})$, between pairs of quantum dots for *ΔF* > 0 (black open circles) and *ΔF* < 0 (red open squares). An apparent inflation in the scatter of strain values is observed at smaller separations, particularly for $\acute{L}(F_{\text{init}})$ ≲2000 nm. (C) Average strain (%$\bar{\varepsilon}$) as a function of $\acute{L}(F_{\text{init}})$, calculated by iteratively excluding data points with the smallest values of $\acute{L}(F_{\text{init}})$at each step. Shaded bands represent the 95% confidence intervals of the mean. (D) Standard error of the mean (SEM) of the average strain values shown in panel (C) plotted as a function of $\acute{L}(F_{\text{init}})$. The minimum SEM occurs at $\acute{L}(F_{\text{init}})$values of 2597 nm and 1699 nm for *ΔF* > 0 (black filled circles) and *ΔF* < 0 (red filled squares), respectively.

**Figure S3:**

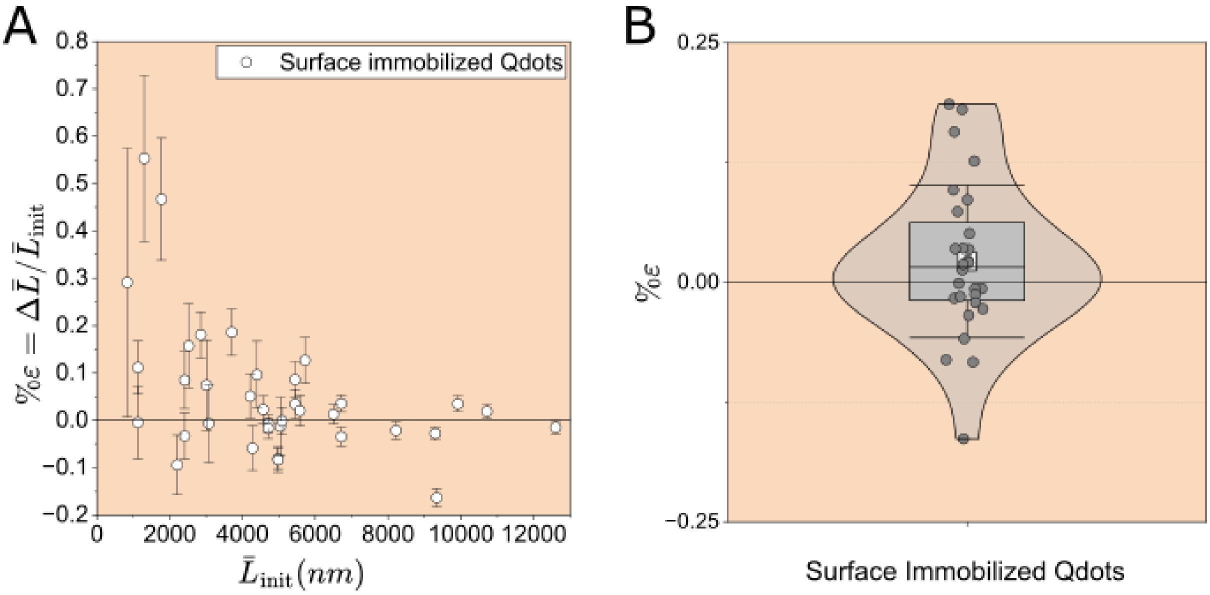


**Relative changes in distances between surface immobilized QDs.** (A) Strain (%ε) of surface-immobilized QDs plotted as a function of the average initial separation, $\acute{L}_{\text{init}}$, between pairs of quantum dots for two different stage positions, shifted by approximately 100 nm along the x-axis. Error bars were calculated using standard error–propagation formulas based on the super-resolution positional tracking uncertainty of each quantum dot. (B) Violin–box plot of strain (%ε) for pairs of quantum dots with $\acute{L}_{\text{init}}$ ≥2500 nm. The mean strain is not statistically different from zero. Error bar is the standard deviation.

**Fig. S4:**

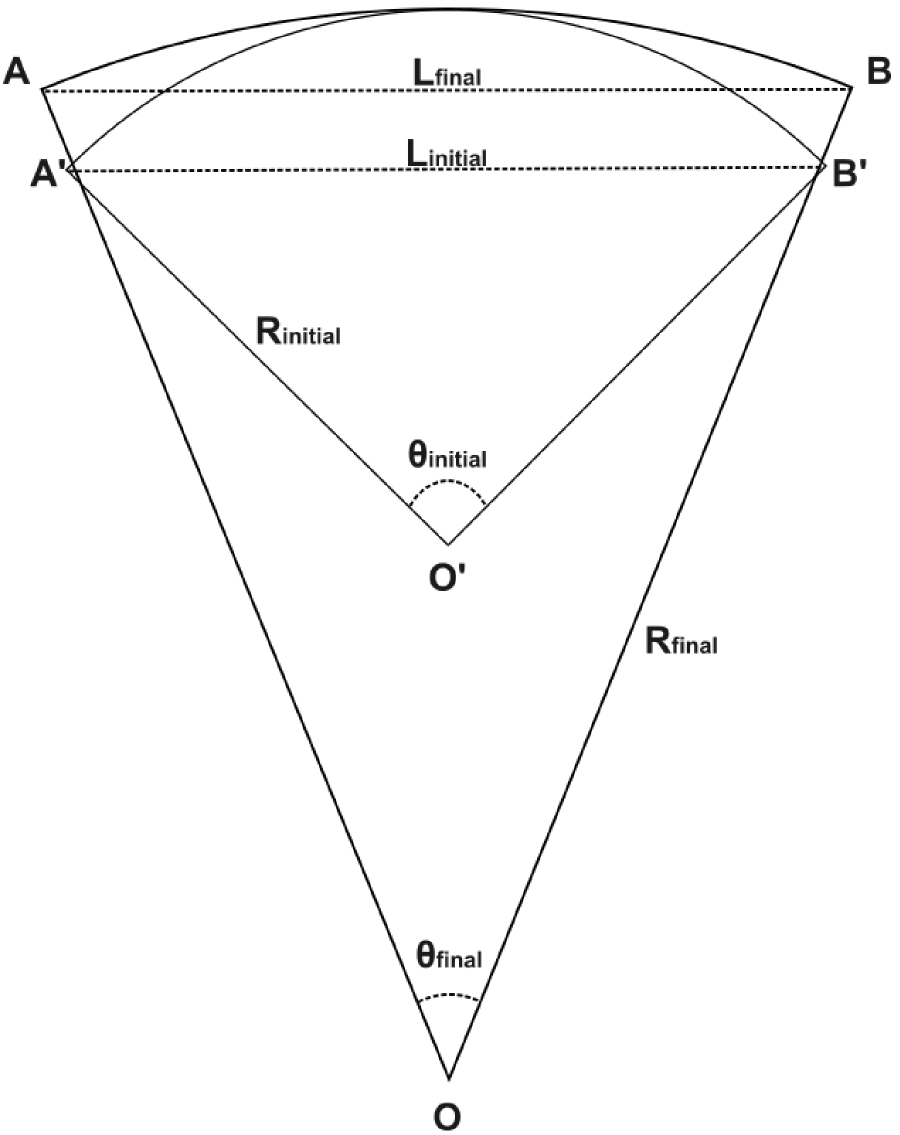


**Relative change in chord length for equal arc length at two different radii of curvature.** Geometric drawing of the change in chord length from $L_{initial}$ to $L_{final}$ upon change of radius of curvature from $1/R_{initial}$ to $1/R_{final}$ at constant arc length, i.e., $R_{initial} \cdot \sin\theta_{initial} = R_{final} \cdot \sin\theta_{final}$.

**Figure S5:**

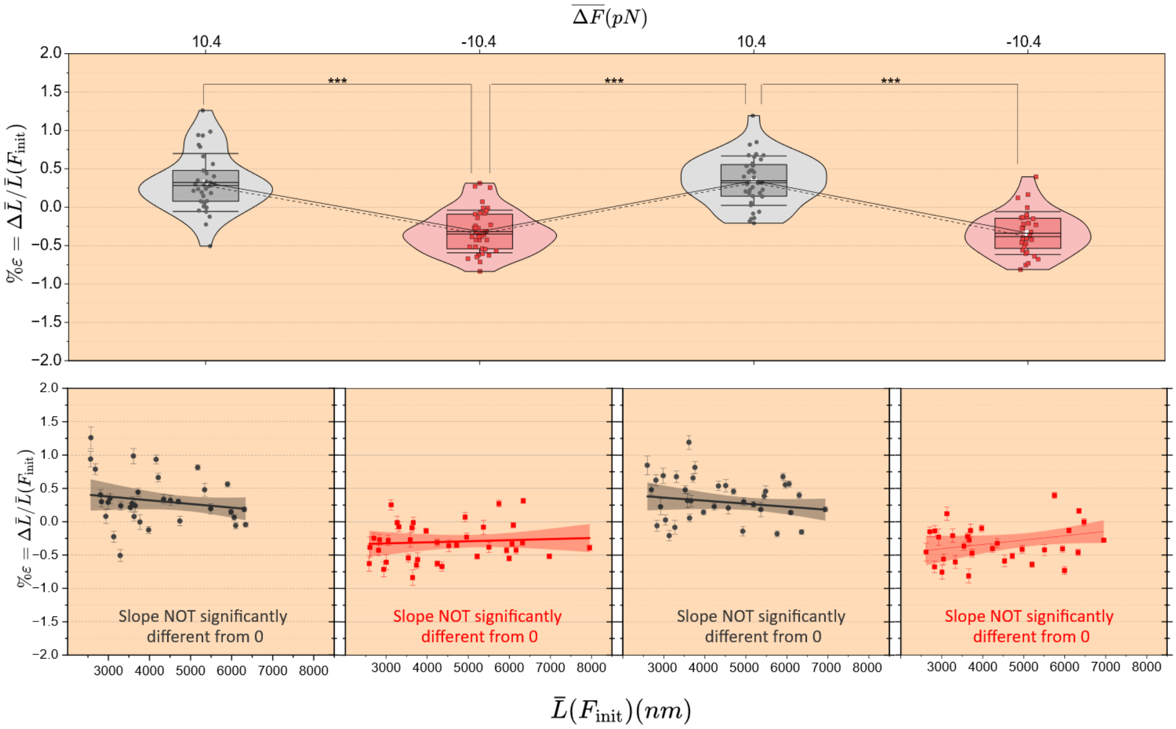


**Strain is independent of the initial distance between QD pairs.** Upper panel: Violin–box plot for the distributions of strain (%ε) for pairs of quantum dots during successive cycles of an average tensile force change of 10.6 pN (same data as shown in the left panel of Fig. 1D). Lower panels: Each plot shows strain (%ε) as a function of the initial separation between pairs of quantum dots and corresponds directly to the violin–box plot distribution displayed above it in the upper panel (number of points in each plot N = 34, 39, 37, 32). Solid lines indicate linear fits, and shaded regions represent the 95% confidence intervals of the fits. In all cases, the fitted slopes were not statistically different from zero, as indicated by the 95% confidence intervals. Error bars for strain were calculated using standard error–propagation formulas based on the super-resolution positional tracking uncertainty of each quantum dot.

**Figure S6:**

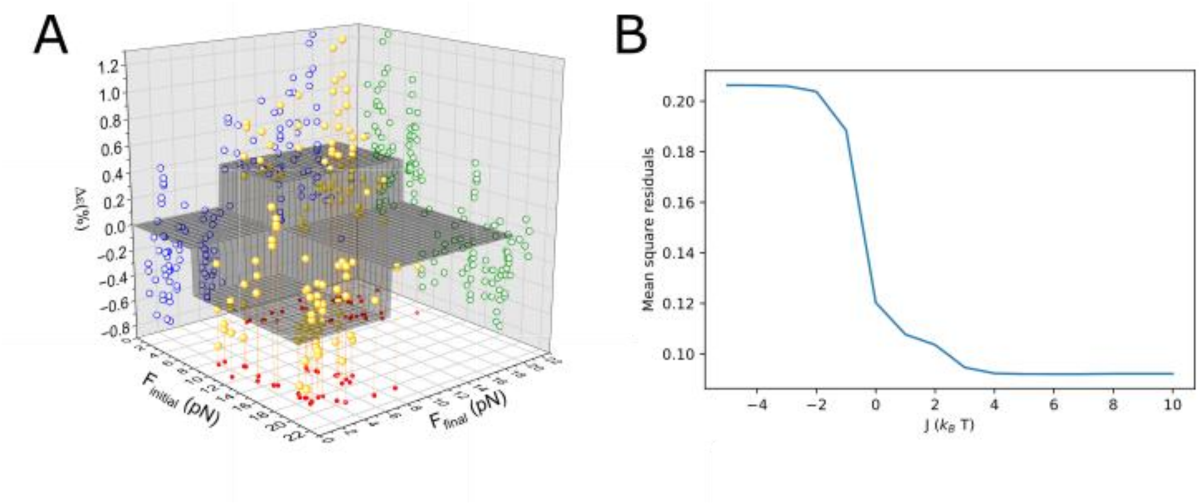


**Three-dimensional fit of ε as a function of F_initial and F_final based on a bistable tubulin lattice and the corresponding Ising model.** (A) Scatter plot of strain $\Delta\varepsilon$ (%) as a function of $F_{initial}$ and $F_{final}$ (yellow 3D spheres, N = 142). The semi-transparent gray surface represents a fit to the data based on the 1D Ising chain model and $\Delta\varepsilon = \varepsilon(\Delta G, F_{final}, J) - \varepsilon(\Delta G, F_{initial}, J)$ (Eq. [3]; see text). Blue and green open circles represent projections of the data onto the $\varepsilon \times F_{final}$ and $\varepsilon \times F_{iitial}$ planes, respectively. The step-like behavior of ε is most clearly apparent in both projections. Red dots represent the projection of the data onto the $F_{final} \times F_{initial}$ plane. (B) Fit residuals as a function of the coupling energy J (Eq. [3]), obtained by fixing J at each value shown while optimizing all other free parameters. The residuals plateau for J ≳ 3 $k_BT$, confirming that the data are insensitive to the precise value of J above this threshold, while the significant increase in residuals for J < 3 $k_BT$ establishes a lower bound on the coupling energy.

**Figure S7:**

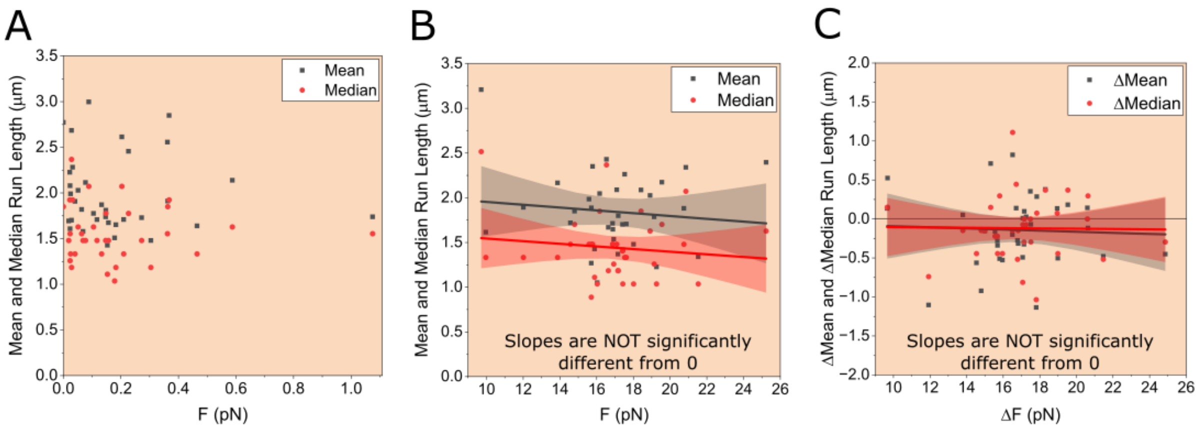


**Mean or median run length of K560 is independent of the tensile force.** (A) Mean (black symbols) and median (red symbols) run lengths of K560 measured on individual microtubules plotted as a function of the initial applied force. (B) Mean and median run lengths of K560 for the same microtubules plotted as a function of the final applied force. (C) Differences in mean and median run lengths calculated from panels (A) and (B) plotted as a function of the corresponding change in force. Solid lines in (B) and (C) indicate linear fits, and shaded regions represent the 95% confidence intervals of the fits.

**Figure S8:**

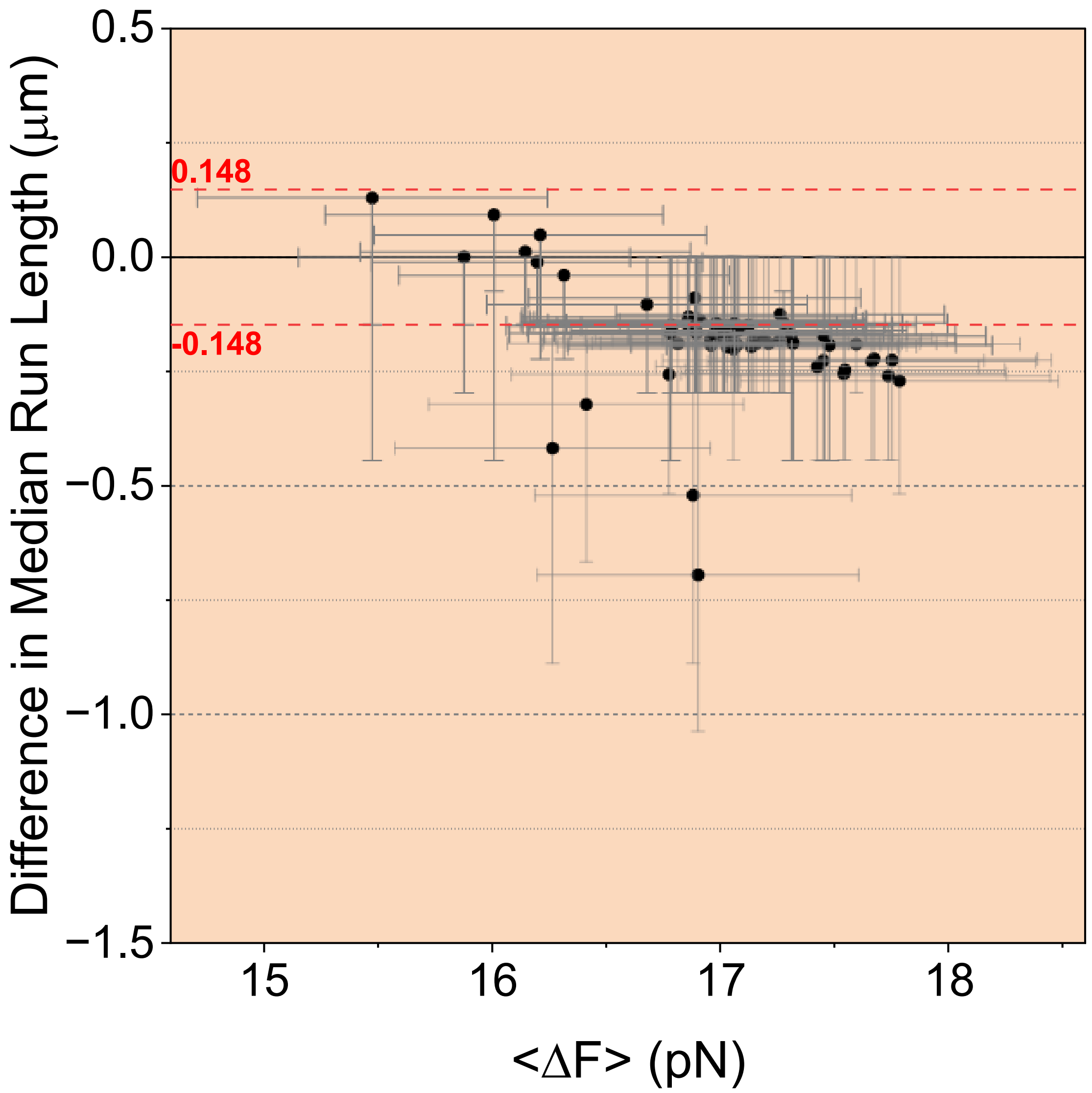


**Differences in median run length of K560 as a function of change in tensile force.** Differences in median run length of hKIF5B(560) as a function of the average change in tensile force for the different subsets shown in Fig. 2E. Error bars represent 95% confidence intervals calculated from 1000 bootstrap replicates. The horizontal red dashed lines indicate the pixel size of the camera (0.148 µm).

**Figure S9:**

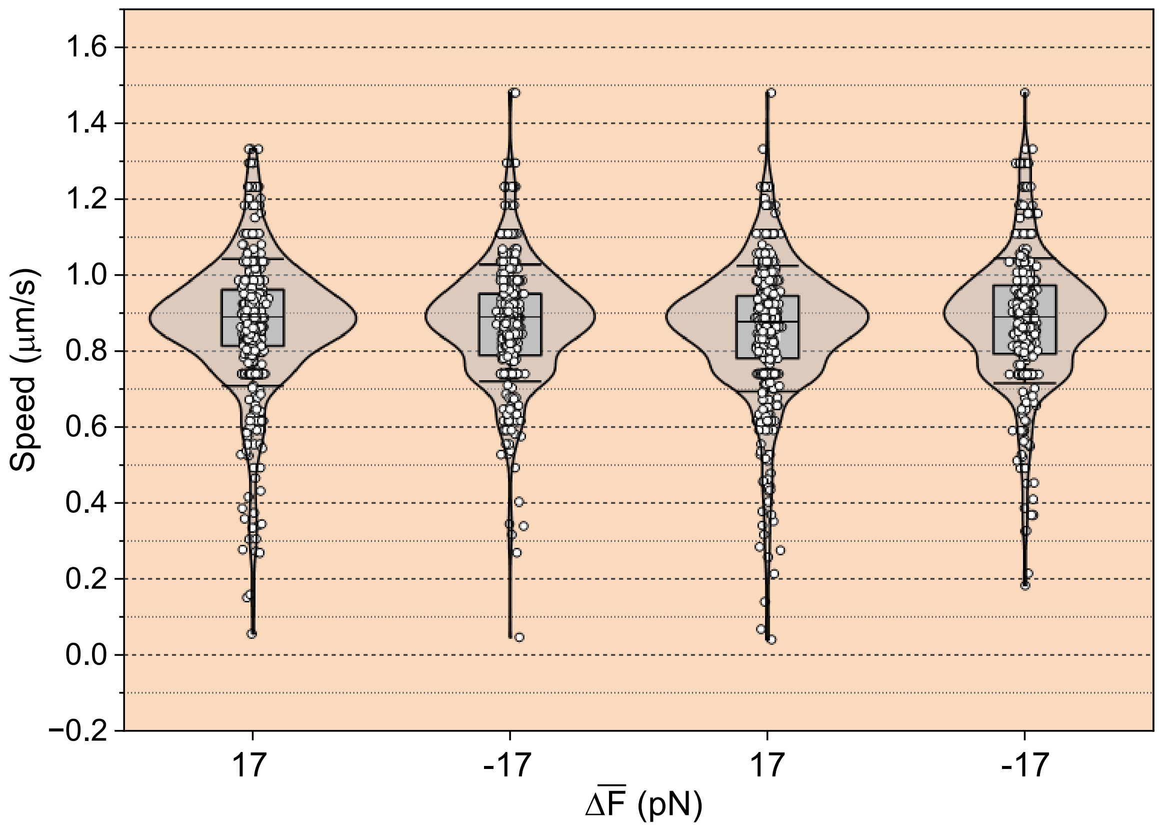


**Speed of K560.** Violin–box plots showing the distributions of hKIF5B(K560) velocities during successive cycles of an average tensile force change of 17 pN. Error bars represent the standard deviation.

**Figure S10:.**

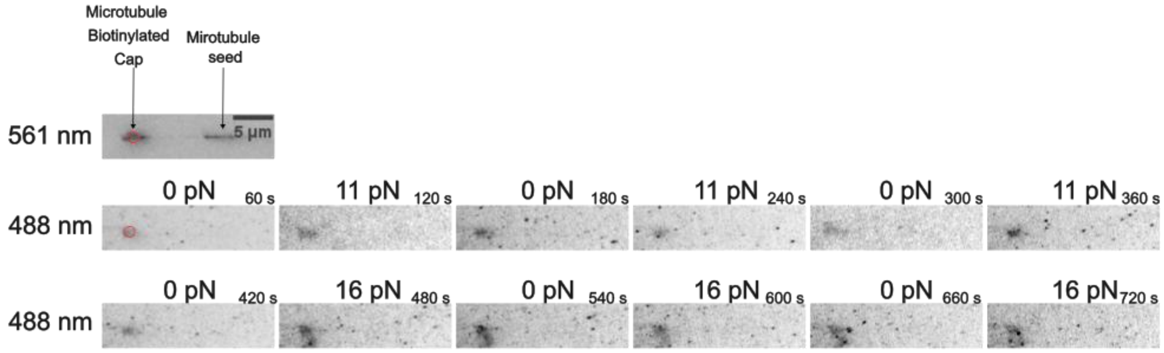


**No incorporation of soluble fluorescent tubulin was observed in the microtubule lattice upon repeated application of tensile force.** First row: The surface-attached microtubule, anchored via its seed, is shown in the Rhodamine/TRITC channel (561 nm). The biotinylated cap, the laser-trapped bead (red circle), and the seed are indicated by vertical black arrows. Second and third rows: Maximum Z-projections in the Alexa 488 channel of the same region shown in the 561 nm channel, displayed over 6 consecutive force cycles in which the force alternates every minute between zero and a non-zero value, as indicated above each sub-panel. The experiment was performed in the presence of 22 nM soluble tubulin (1 mM GTP), of which 2% was fluorescently labelled with Alexa 488. This concentration is at least 6-fold above the dissociation constant of the αβ-tubulin heterodimer, $K_d$ = 3.69 ± 0.65 nM ensuring that tubulin is predominantly in the dimeric state under these conditions (Fineberg et al., J. Mol. Biol. 2020). The trapped bead carries attached TRITC-labelled microtubule fragments, resulting in a locally high TRITC concentration and consequent bleedthrough into the 488 nm imaging channel. In addition, the bead diameter (2r = 784 nm) is comparable to the 488 nm excitation wavelength, making it a highly efficient Mie scatterer at this wavelength. Both effects contribute to the bright glare observed in the 488 nm channel at the bead position.

**Figure S11:**

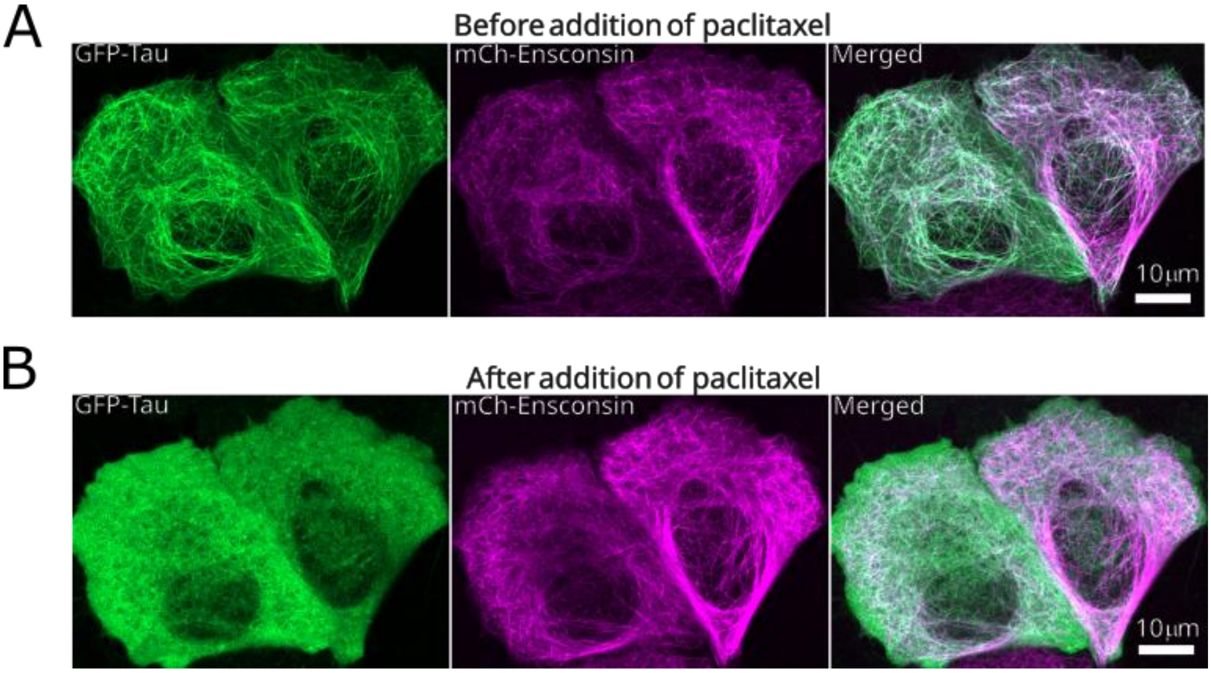


**Paclitaxel treatment of HeLaM cells overexpressing mCh-enscosnin and eGFP-tau.** Fluorescence images of HeLaM cells co-expressing eGFP-tau and mCh-ensconsin, shown individually and as a merged overlay, in **(A)** the absence and **(B)** the presence of 10 μM paclitaxel. Upon paclitaxel addition, eGFP-Tau is displaced from the microtubule lattice and redistributes to the cytosol, while mCh-Ensconsin remains associated with microtubules.

**Figure S12:**

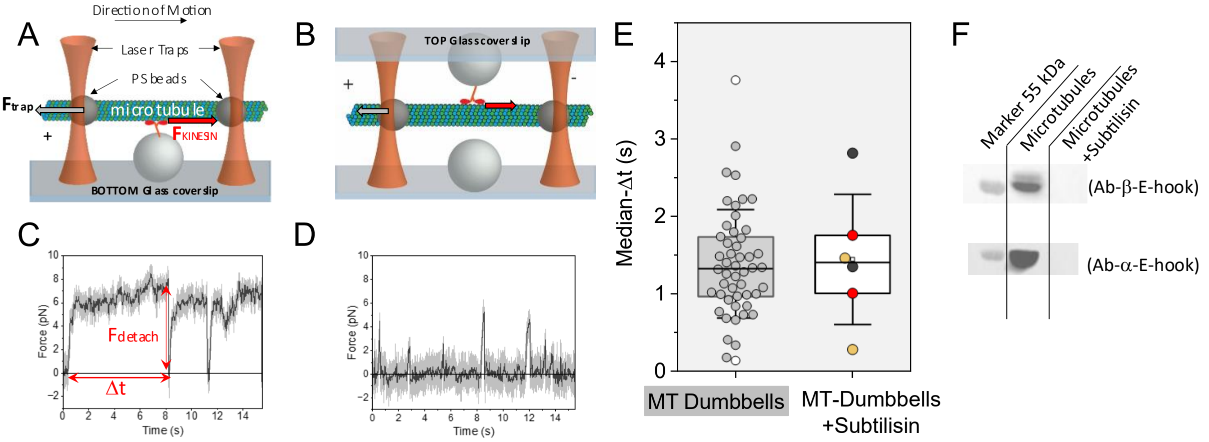


**Heterogeneity of microtubule–hKIF5B(560) interaction under opposing load is preserved following subtilisin cleavage of the tubulin C-terminal tails.** (A–B) Schematic depictions of the three-bead assay in which single kinesin-1 molecules anchored on surface-immobilised spherical pedestals, interact with the same microtubule at different locations — bottom (A) and top (B) — in the presence of 1 mM ATP. (C–D) Representative force traces recorded at the bottom (C) and top (D) locations of the same microtubule. The detachment force $F_{detach}$ and the attachment duration Δt for each force ramp are indicated by red double-headed arrows. (*E*) Box-plot with overlaid scatter of median Δt values measured at top and bottom locations for untreated (grey scatter points, with the two extreme values in (C) and (D) traces shown as white points; n = 25 microtubules) and subtilisin-treated (coloured scatter points; n = 3 microtubules) GDP-paclitaxel microtubules. Data for untreated microtubules are from a previously published study (1, 15). For subtilisin-treated microtubules, top and bottom location measurements from the same microtubule are shown in the same colour, with each microtubule represented by a distinct colour. (F) Western blot of microtubule samples before and after subtilisin treatment, confirming complete removal of both α- and β-tubulin C-terminal tails. Subtilisin treatment was performed as previously described (16).
